# Age-related decline of synaptic plasticity is regulated by neuro-androgen and neuro-estrogen in normal aging of hippocampus

**DOI:** 10.1101/2025.08.17.670749

**Authors:** Suguru Kawato, Yasushi Hojo, Mari Ogiue-Ikeda, Mika Soma, Minoru Saito, Jonghyuk Kim, Arisa Munetomo, Shimpei Higo, Hirotaka Ishii, Asami Kato, Hideo Mukai, Ayako Hiragushi, Takuji Shirasawa, Takahiko Shimizu, Shigeo Horie

**Affiliations:** Department of Biophysics and Life Sciences, Graduate School of Arts and Sciences, The Univ. Tokyo, Tokyo, Japan; Department of Cognitive Neuroscience, Faculty of Pharma-Science, Teikyo Univ., Tokyo, Japan; Department of Urology, Graduate School of Medicine, Juntendo University, Tokyo, Japan; Department of Biochemistry, Saitama Medical University, Saitama, Japan.; Department of Biosciences, College of Humanities and Sciences, Nihon University, Tokyo, Japan; Molecular Gerontology, Tokyo Metropolitan Institute of Gerontology, Tokyo, Japan.; Department of Aging Control Medicine, Graduate School of Medicine, Juntendo University, Tokyo, Japan; Department of Mechanism of Aging, National Center for Geriatrics and Gerontology, Aichi, Japan

**Keywords:** aging, hippocampus, neuroandrogen, neuroestrogen, spine

## Abstract

We revealed a good relationship between age-dependent decrease in the hippocampal dendritic spine density and age-dependent decrease in hippocampal androgen and estrogen levels with normal aging of male rats.

Approximately 25% decrease in the spine density was observed in hippocampal CA1 region by going from 3 month-old (3m; young adult) to 24 month-old (24m; aged). We found a significant age-induced decrease in hippocampal neuro-androgen levels by going from 3m to 24 m using mass-spectrometric analysis. The hippocampal levels of testosterone (T) and dihydrotestosterone (DHT) dramatically decreased from 17 nM T and 7 nM DHT at 3 m to 17/100 nM T and 7/15 nM DHT at 24 m. On the other hand, hippocampal estradiol (E2) was moderately decreased with aging, from 8 nM at 3 m to 2 nM at 24m.

Comprehensive analysis of mRNAs of hippocampal steroidogenic enzymes and receptors showed an age-dependent decrease in their expression levels by approximately 50% (P450(17α)), 25% (17ꞵ-hydroxysteroid dehydrogenase) and 0% (5α-reductase and P450arom). Androgen receptor AR was moderately decreased but estrogen receptor ERα was not decreased with aging.

The 25% decrease in the spine density with aging may be due to a balance between considerably decreased T and DHT levels (spine decrease factor) and remained moderately high E2 level (spine increase factor) in the 24m hippocampus. Aged hippocampus still has moderate capacity of sex-steroid synthesis and their functions.

Interestingly, DHT-supplementation and T-supplementation recovered the spine density at 24m.

## 1. Introduction

Our increasing life span is expected to decrease quality of life due to age-related declines in cognitive functions, due to not only pathological changes such as Alzheimer’s disease (AD) but also normal aging.

In healthy humans, many literatures have described age-related perturbations in hippocampal activity coincident with deficits of learning and memory (Daselaar et al 2006) (Dennis et al 2008) (Beeri et al 2011).

Similarly, rodent models of normal aging demonstrate strong correlations between impaired performance of aged rats on behavioral tests of hippocampus-dependent learning and memory and decreased hippocampal neuron activity. The performance of spatial learning (dependent on CA1 region) and contextual fear learning (dependent on CA3 region) decline along with aging (Barnes 1987) (Burke & Barnes 2006) (Veng et al 2003). Age-dependent impairment of electrophysiological properties of hippocampal function is related to unstable encoding of spatial representations (Barnes et al 1997) (Kumar et al 2007) (Norris et al 1996) (Rosenzweig & Barnes 2003) (Wilson et al 2003).

Aging-dependent pathological cognitive decline such as AD is a result of neuronal cell death or cell loss (Morrison & Hof 1997) (Morrison & Hof 2002) (West et al 1994). In case of normal aging, however, the number of neurons in the hippocampus and the cerebral cortex (CC) do not change along with aging (Burke & Barnes 2006) (Kelly et al 2006) (Morrison & Hof 1997) (Morrison & Hof 2002) (Rapp & Gallagher 1996) (Rasmussen et al 1996) (West et al 2004).

These results suggest that the decline in the learning and memory with normal aging is likely to occur at a synaptic level, because of no neuron loss. Dendritic spines (post-synapses) in the hippocampus correlate with the performance of learning and memory. Increase in the density and the size of spines is associated with learning and memory tasks (Moser et al 1994, Moser et al 1997, O’Malley et al 2000). Structural synaptic plasticity is so important to study, because based on mounting evidence, it appears that growth of spines and formation of their synapses represent a morphological substrate for learning and memory (Kasai et al., 2003; Lang et al.,2004; Silva, 2003).

We focus on sex steroids as strong candidates for neurotrophic factors that correlate with aging effects on decline of the cognitive function in addition to reproductive system (Midzak et al 2009) (Chen et al 1994).

Decline in sex steroid, especially 17β-estradiol (E2), has been considered as an important factor which is involved in age-related female neural dysfunction, since E2 significantly decreases in plasma upon menopause that elicits cognitive decline and dysfunction of Hypothalamus-Pituitary-Gonadal axis (Kawakita et al 2021) (Kim et al 2017). The relationship between neural function and E2 has been extensively investigated in the male and female rodent hippocampus, including the modulatory effect of E2 on synaptic plasticity in CA1 region (Bi et al 2000) (Foy et al 1999) (Hasegawa et al 2015) (Mukai et al 2007) (Mukai et al 2010) (Ooishi et al 2012a) (Ooishi et al 2012b) (Vouimba et al 2000). The beneficial effects on dendritic spines by E2 replacement therapy have been investigated in aged female rodents and nonhuman primates (Hao et al 2007) (Hara et al 2015) (Morrison & Baxter 2012).

The mechanisms of androgen effects on synaptic plasticity have also attracted much attention. For example, perfusion of testosterone (T) and dihydrotestosterone (DHT) significantly increased dendritic spine density within 2 h in rat hippocampus slices (Hatanaka et al 2015). These rapid spine increases were driven by kinases via membrane synaptic AR receptor. T administration increases synaptic density in the dentate gyrus of aged mice (Fattoretti et al 2019). Castration of young male rats decreased serum T, and T replacement improved spatial memory (Wagner et al 2018). T replacement also improved spatial memory in aged male rats whose serum T level is low (Jaeger et al 2020). SAMP8 mouse is a senescence accelerated mouse which exhibits cognitive impairment and hypogonadism. Investigations of male SAMP8 mice showed the relationship between plasma T decline and age-related neural dysfunction (Ota et al 2012). Interestingly, T and DHT replacement in male SAMP8 inhibited cognitive decline through an increase of sirtuin 1 protein (SIRT1) expression (Ota et al 2012). In relation to AD, apparent relationship between decrease in T level and progression of AD was observed in both human and rodents model of AD (Rosario et al 2004) (Rosario et al 2010).

Human data also indicate a significant decline in plasma T of men due to chronological aging (Harman et al 2001) (Morley et al 1997) (Travison et al 2007). Plasma T declines across the men’s life span represents an issue of great concern for cognitive impairment. In many cases, the aging effects had been attributed to the decline in plasma T and E2, which are synthesized in testis and ovary (Janmaat et al 2011, Wu et al 2009). However, the hippocampus is able to locally synthesize sex steroids *de novo* in addition to the target of sex steroids (Fig. 1) (Hojo et al 2004) (Hojo et al 2008) (Kato et al 2013) (Kawato et al 2002) (Kimoto et al 2010).

**Figure 1:**
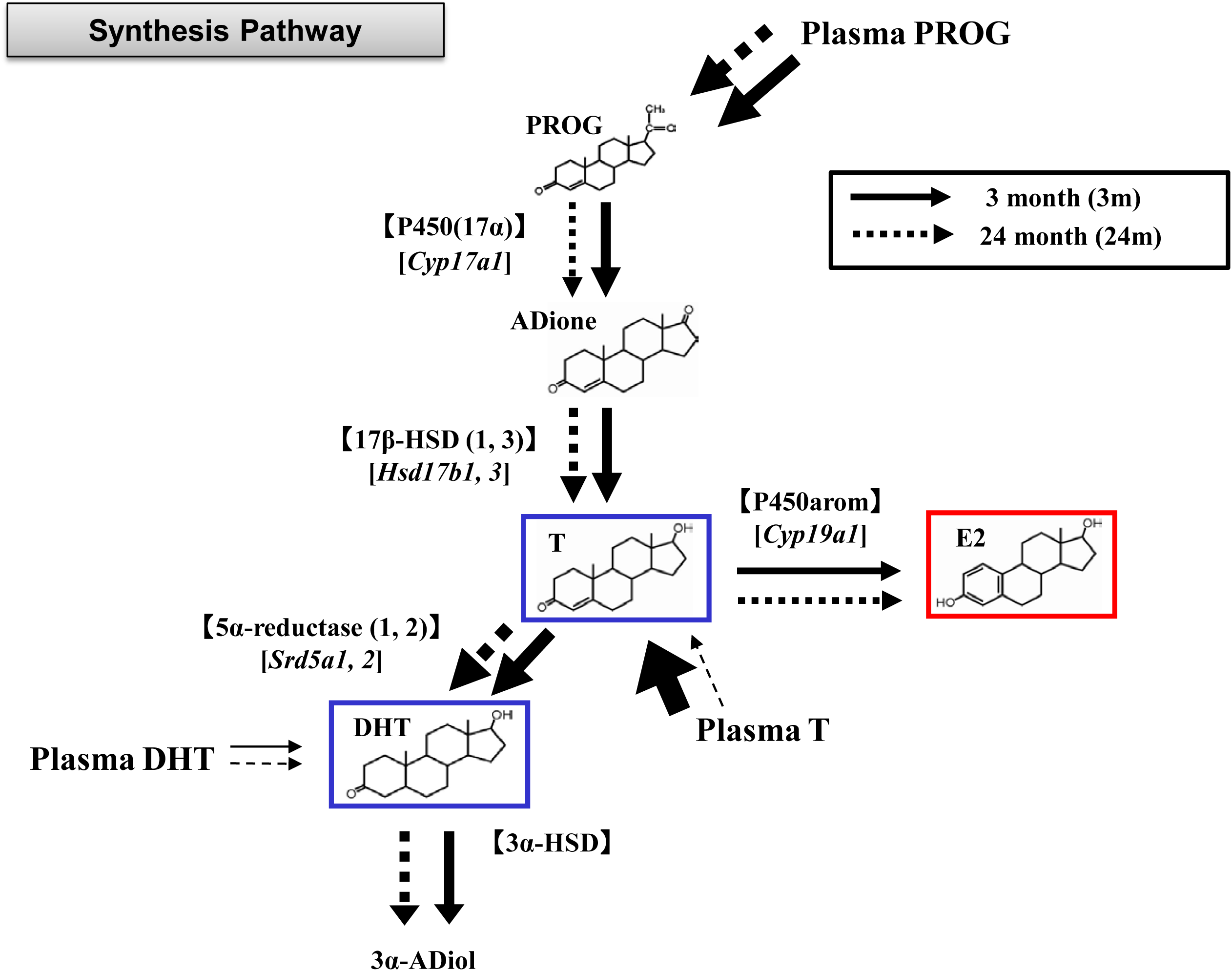
Male hippocampal steroid metabolism pathways, including enzymes and receptors examined in the current study. Straight arrows indicate 3m hippocampal pathways and dotted arrows indicate 24m hippocampal pathways. Relative thickness of arrow implies flow rate of metabolism. Sufficient PROG is supplied to the hippocampus via blood PROG circulation in both 3m and 24m. Hippocampal androgen synthesis capacity from PROG to T depending on P450(17α) and 17β-HSD (1,3) decreased at 24m. Because T is rapidly and efficiently converted to DHT by 5a-reductase (highly expressed even at 24m), resultant T level may be low (∼0.15 nM). T supply from circulation is very poor (∼0.3 nM plasma) at 24m. However at 3m, 70% of hippocampal T (15 nM) is supplied via blood circulation (Hojo et al 2009). T is converted to E2 by P450arom, rather slowly. E2 is very stably present without degradation (Hojo et al 2009), but DHT is further converted to 5α-androstanediol by 3α-hydroxysteroid dehydrogenase (3α-HSD). Very low plasma T, DHT levels at 24m cannot contribute to hippocampal T, DHT levels. Note that plasma E2 level is already very low at 3m and 24 male. PROG: progesterone, ADione: androstenedione, T: testosterone, DHT: dihydrotestosterone, E2: estradiol, 3α-ADiol: 3α-androstanediol.

Evidence of the role of hippocampus synthesized sex steroids in hippocampal function has been accumulated, including neurogenesis (Okamoto et al 2012), LTP (Di Mauro et al 2015, Di Mauro et al 2017, Grassi et al 2011) and memory consolidation (Tuscher et al 2016).

The purpose of the current study is, therefore, to demonstrate the relationship between age-related decrease in dendritic spines and aging-dependent decline in hippocampal sex steroidogenic systems. These investigations focus on hippocampal levels of androgen and estrogen as well as their receptors. We observed in the hippocampus, a good correlation between the aging-dependent decrease in the density of CA1 spines and the considerable decrease in androgen levels that reduced to ∼1/100 fold by going from 3 month-old (3m, young adult) to 24 month-old (24m, old). The decrease was also observed in mRNA expression of androgen receptor (AR) and some androgen-synthesizing enzymes. The significant recovery of the spine density with androgen replacement at 24m hippocampus supports the capacity of hippocampal androgen that can prevent synaptic impairment with aging. Interestingly, the 24m hippocampal E2 level was only moderately decreased to 2 nM (1/4 fold of 3m hippocampus), and no decrease in P450arom (E2 synthase) and ERα expression was observed, suggesting that E2 can still support synaptic plasticity in the absence of androgen in aged hippocampus.

## 2. Results

### 2.1. Dendritic spines in CA1 pyramidal neurons

#### 2.1.1. Age-induced decrease of spines

We investigated age-induced change in spine density and morphology of the hippocampal CA1 stratum radiatum, for 24m and 3m male rats (Fig. 2A). Spine analysis was performed for secondary branches of the apical dendrites located 100-200 µm away from the pyramidal cell body, in the middle of the stratum radiatum of CA1 region.

**Figure 2:**
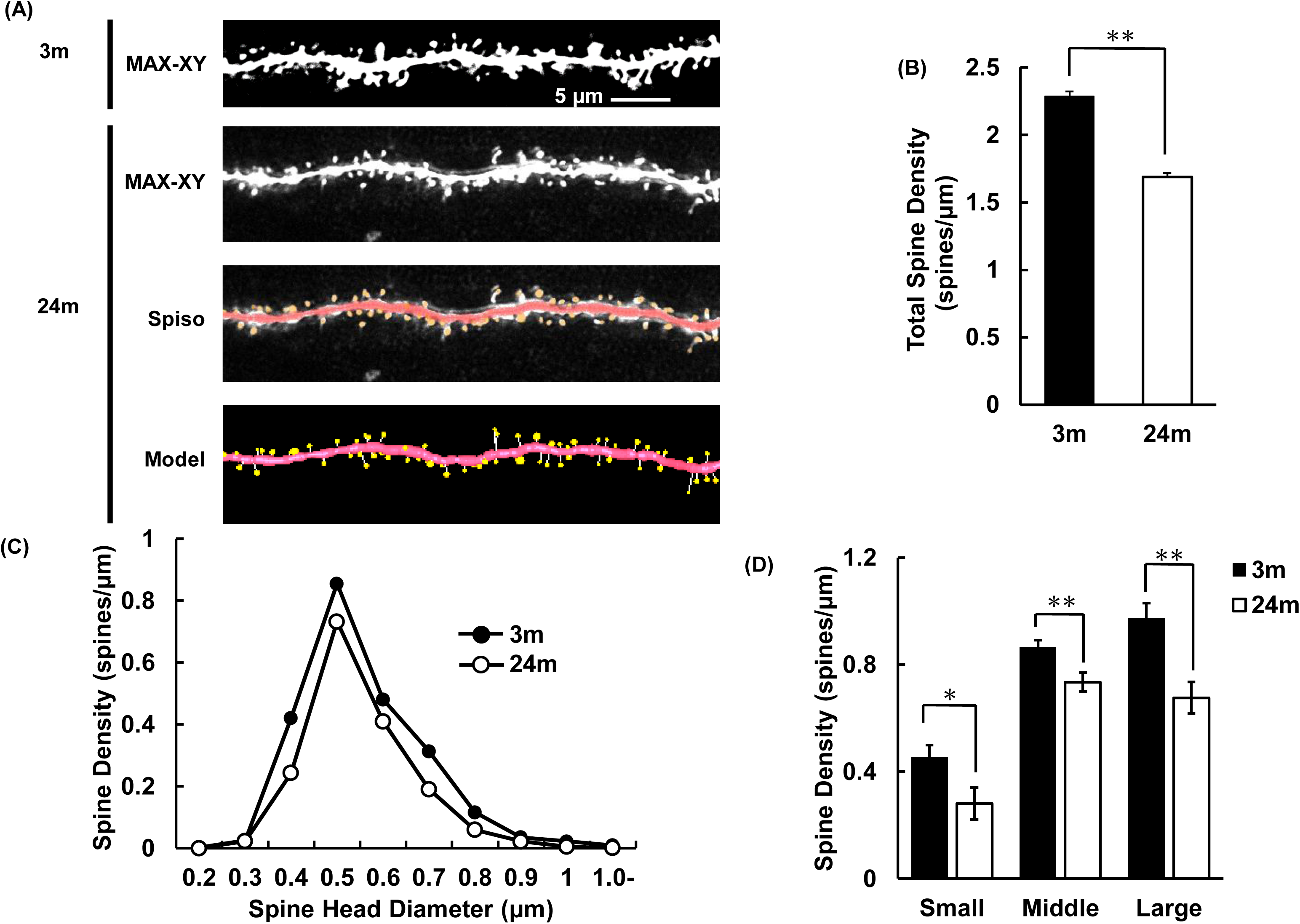
Age-induced decrease of the spine density in hippocampal CA1 pyramidal neurons. Spines were analyzed along the secondary dendrites of CA1 pyramidal neurons in the stratum radiatum in 3m and 24m male rats. (A) Representative images of confocal micrographs; the spines along dendrite in 3m (3m) and 24m (24m). Maximal intensity projection onto XY plane from z-series confocal micrographs (MAX-XY), images analyzed by Spiso-3D (Spiso), and three dimensional model illustrations (Model) are shown together. Scale bar, 5 μm. (B) The total spine density of 3m and 24m. Vertical axis represents the average number of spines per 1 μm of dendrite. Data are represented as mean ± SEM. Closed column, 3m; open column, 24m. (C) Histogram of spine head diameters in 3m (closed circle) and 24m (open circle). (D) Density of three subtypes of spines in 3m (closed column) and 24m (open column). From left to right, small-head spines (Small), middle-head spines (Middle), and large head spines (Large). Data are represented as mean ± SEM. Statistical significance yielded \**p* < 0.05 and \*\**p* < 0.01.

The total spine density showed age-induced decrease, from 2.29 spines/μm (3m) to 1.69 spines/μm (24m) which corresponds to approx. 26% decrease (Fig. 2B).

The morphological changes in spine head diameter were also assessed. Nearly 95 % of spines have clear heads and necks, and we classified the head diameter of spines into three categories: small-head spines (0.2–0.4 μm), middle-head spines (0.4–0.5 μm), and large-head spines (0.5-1.0 μm).

Along with aging, the density of all three types of spines was decreased. The density of large-head spines was decreased from 0.97 spines/μm (3m) to 0.68 spines/μm (24m) (Fig. 2C, 2D). The density of middle-head spines was decreased from 0.86 spines/μm (3m) to 0.73 spines/μm (24m). The density of small-head spines also decreased from 0.45 spines/μm (3m) to 0.28 spines/μm (24m).

#### 2.1.2. Recovery of age-induced spine decrease by androgen administration

Upon single androgen administration for 24m rats, the spine density recovered after 17 h (Fig. 3A).

**Figure 3:**
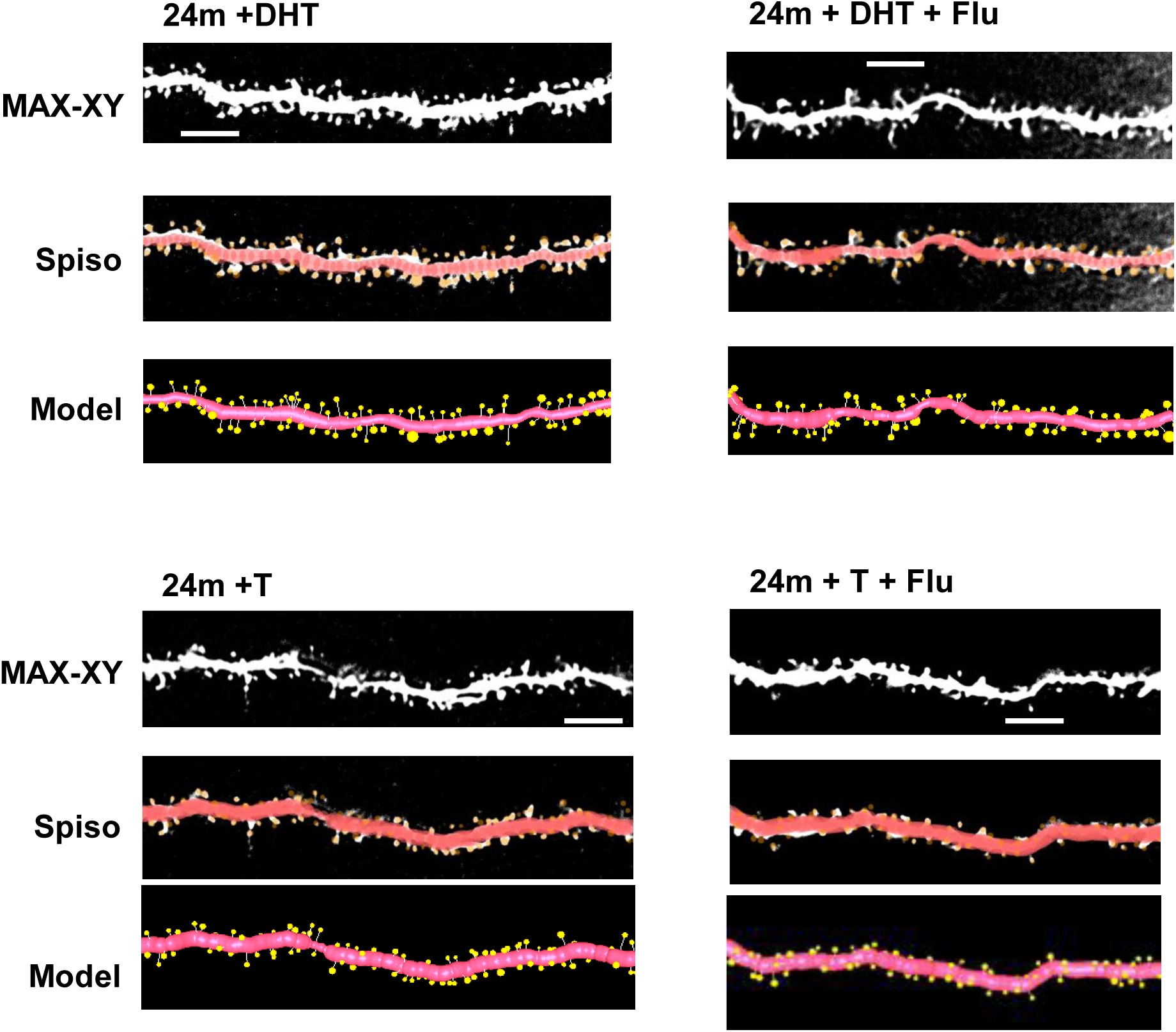
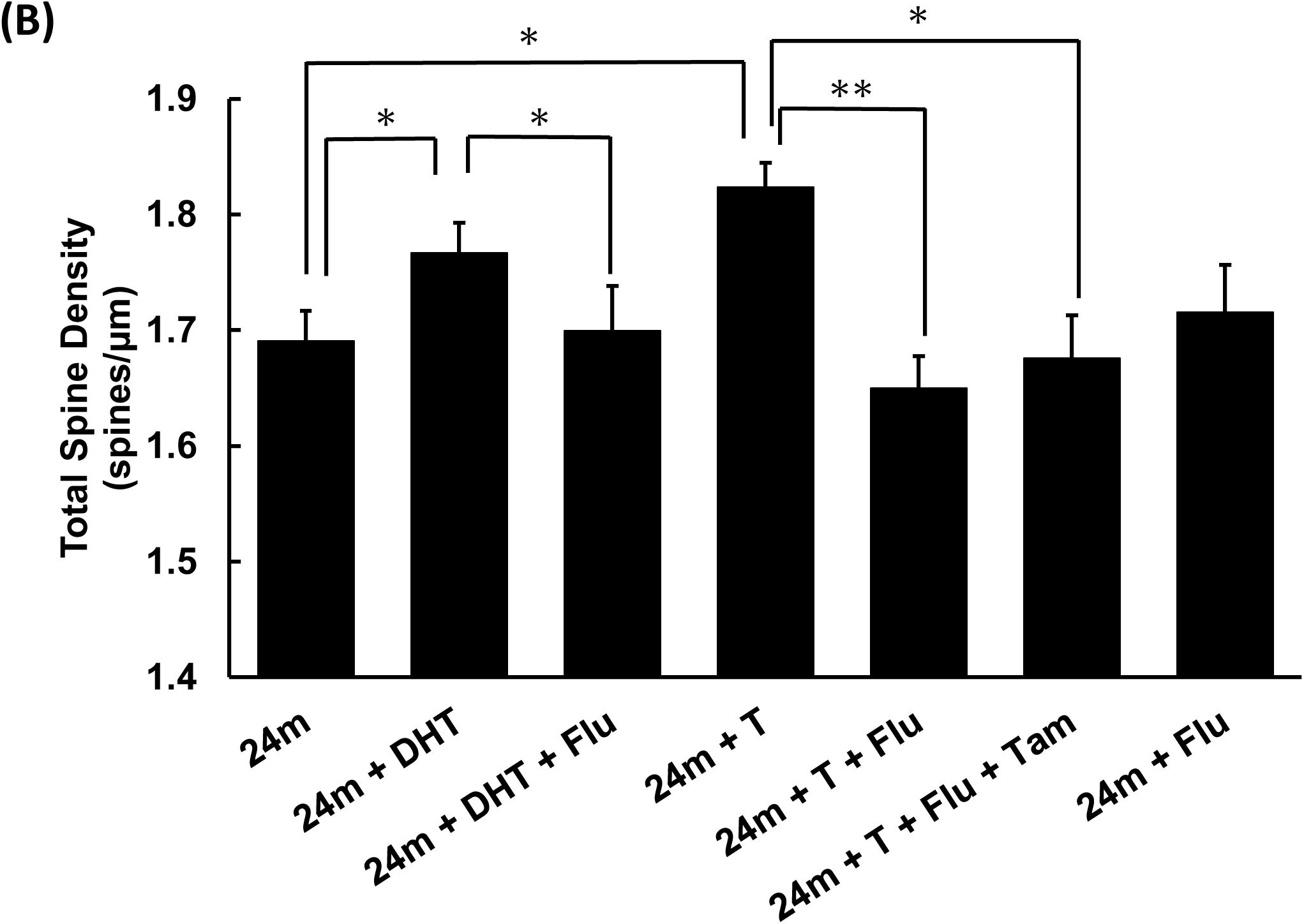
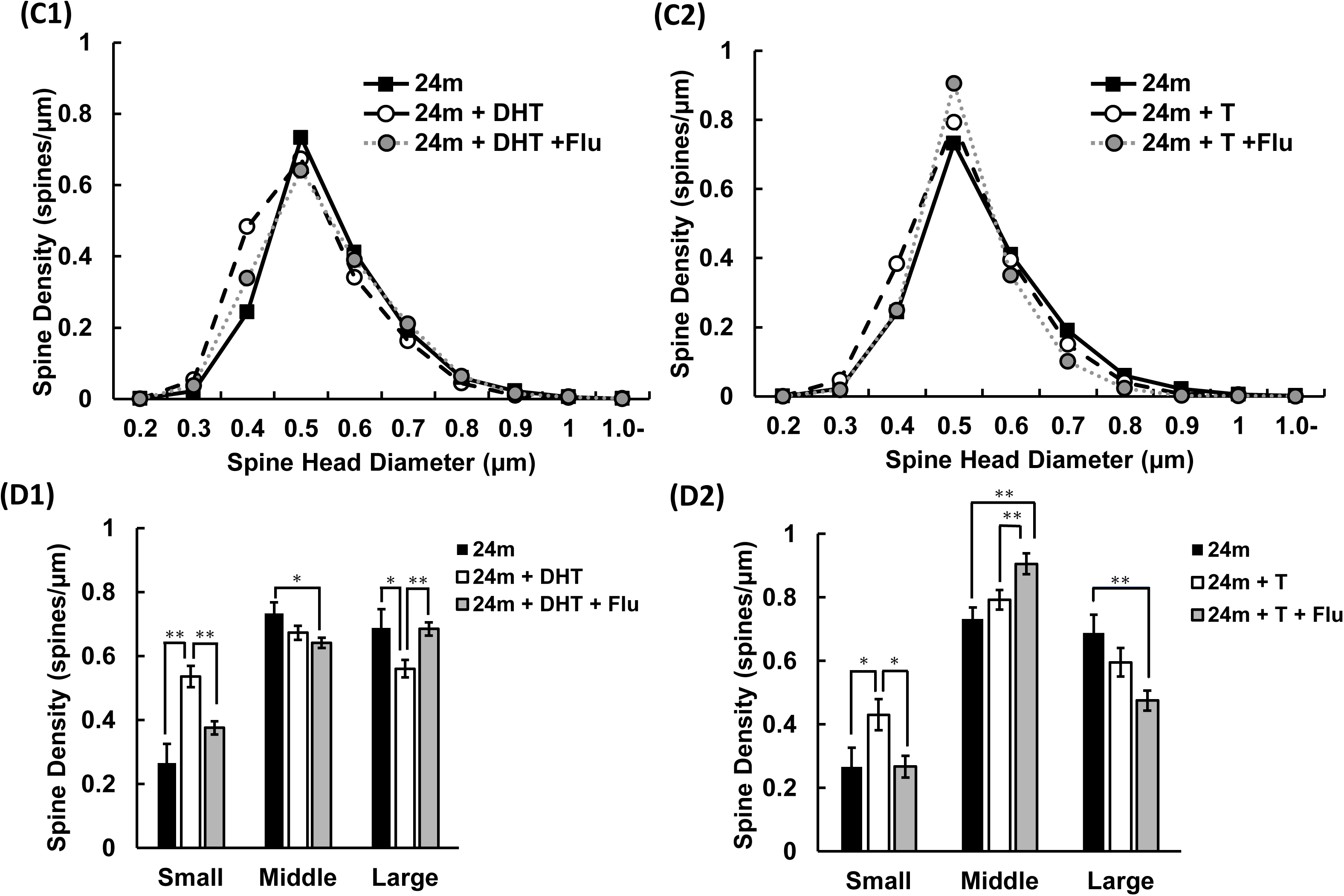
Androgen (DHT and T) - induced increase of the spine density in hippocampal CA1 neurons of 24m male rat, after 17 h of androgen administration. (A) Representative images of confocal micrographs; the spines along dendrite in 24m + DHT, 24m + DHT + Flu, 24m + T, 24m + T + Flu. Maximal intensity projection onto XY plane from z-series confocal micrographs (MAX-XY), images analyzed by Spiso-3D (Spiso), and three dimensional model illustrations (Model) are shown together. Scale bar, 5 μm. (B) Comparison of the total spine density after androgen and blocker administration. (C and D) Morphology change in spines. (C1 and C2) Histogram of spine head diameters in 24m (closed square), 24m + DHT (open circle) and 24m + DHT + Flu (gray circle), and 24m (closed square), 24m + T (open circle), 24m + T + Flu (gray circle). (D1 and D2) Density of three subtypes of spines, small-head spine (Small), middle-head spine (Middle), and large-head spine (Large). From left to right of (D1), 24m (closed column), 24m + DHT (open column), 24m + DHT + Flu (gray column). From left to right of (D2), 24m (closed column), 24m + T (open column), 24m + T + Flu (gray column). Data are represented as mean ± SEM. Statistical significance yielded \**p* < 0.05 and \*\**p* < 0.01.

##### Total spine density analysis

DHT administration (1mg/kg) increased the total density of spines from 1.69 to 1.77 spines/μm (F = 2.984, p = 0.009, two-way ANOVA; p = 0.0172 for 24m vs 24m + DHT, Tukey-Kramer multiple comparison’s test), which corresponds to approx. 5% increase. T administration (1mg/kg) increased the total spine density from 1.69 to 1.82 spines/μm (F = 7.2224, p = 0.008, two-way ANOVA; p = 0.0392 for 24m vs 24m + T, Tukey-Kramer multiple comparison’s test), which corresponds to approx. 8% increase (Fig. 3B). Next, to clarify the involvement of androgen receptor (AR) in the recovery effect of androgen administration, we performed s.c. administration of flutamide (30 mg/kg), a specific antagonist against AR, 1 h before androgen administration. Co-administration of flutamide and DHT decreased the total spine density back to the control level (Fig. 3B). Co-administration of flutamide and T also decreased the total spine density back to the control level (Fig. 3B).

The additional effect of tamoxifen, an antagonist against estrogen receptor, to flutamide was investigated, because a part of T might be converted to estradiol. Simultaneous administration of tamoxifen (1 mg/kg) and flutamide (30 mg/kg) to T-injected group did not further decrease the total spine density of flutamide plus T injected group. As a control, no effect of flutamide alone (without androgen administration) was checked.

##### Spine head diameter analysis

The morphological changes in spine head diameter were also assessed. Upon DHT supplementation, small-head spines were significantly increased from 0.27 to 0.54 spines/μm (p < 0.0001 for small-head, Tukey-Kramer), while some decrease in large-head spines was observed (Figs. 3C1, 3D1). Upon T supplementation, small-head spines were mainly increased from 0.27 to 0.43 spines/μm (p = 0.0468 for small-head, Tukey-Kramer), without changes in middle-head and large-head spines (Figs. 3C2, 3D2).

Co-administration of flutamide (Flu) and DHT blocked the DHT-induced increase of small-head spines and the DHT-induced decrease of large-head spines, without changing middle-head spines (p < 0.0001 for small-head, p = 0.0002 for large-head, Tukey-Kramer) (Figs. 3C1, 3D1). Co-administration of Flu and T blocked the T-induced increase of small-head spines (p = 0.0273 for small-head, Tukey-Kramer) (Figs. 3C2, 3D2).

### 2.2. Age-induced decrease in androgen and estrogen levels in the hippocampus from 3m to 12m and 24m

Mass-spectrometric analysis with LC-MS/MS was performed to accurately determine the change in levels of T, DHT and E2 with aging (Fig. 4). The concentrations of T, DHT and E2 were determined by a chromatogram analysis of the fragmented ions of T-17-picolinoyl-ester, DHT-17-picolinoyl-ester and E2-3-pentafluorobenzyl (PFBz)-17-picolinoyl-ester, as shown in Supplementary figures (Fig. S1).

**Figure 4:**
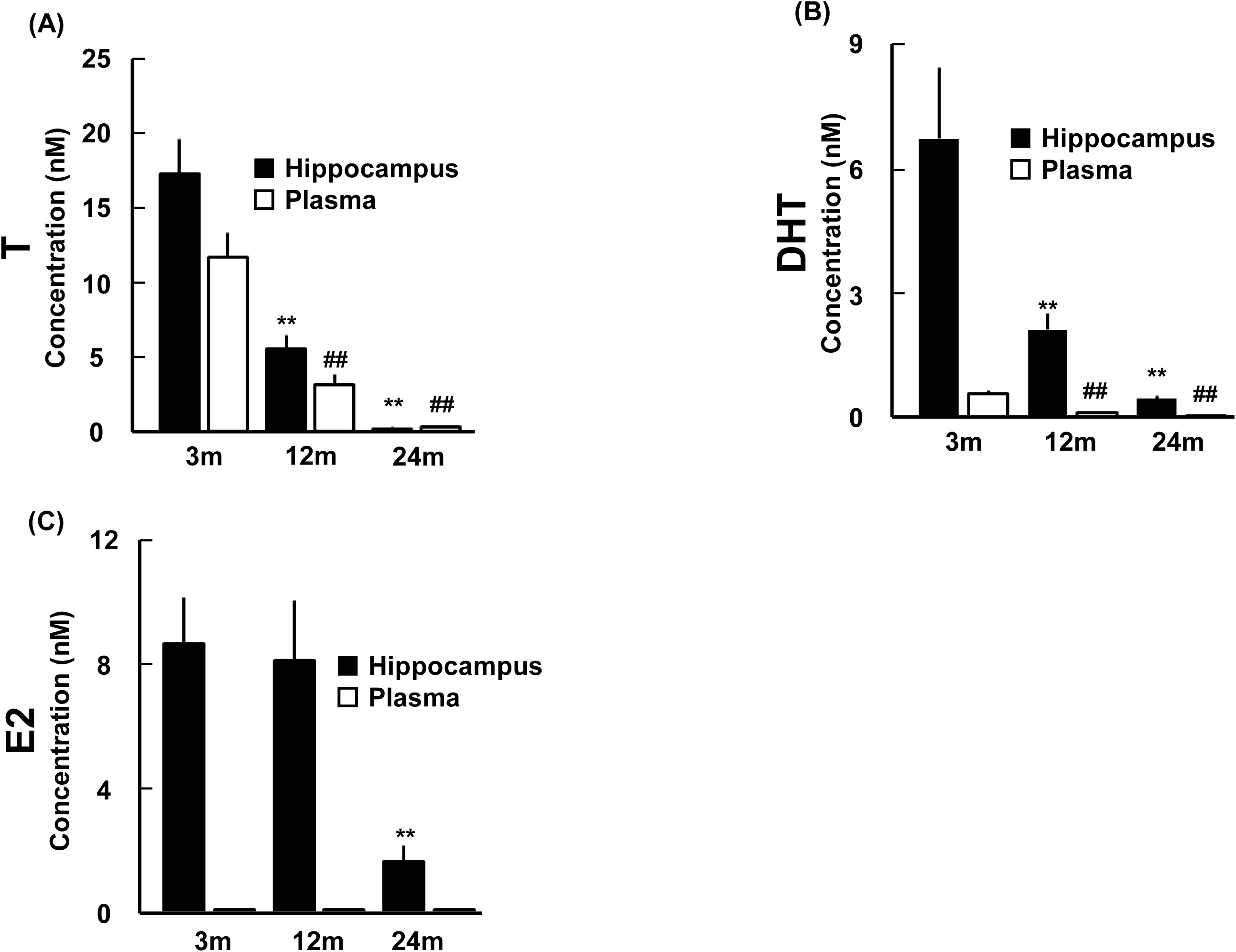
Age-induced decrease in T, DHT and E2 levels in the hippocampus and plasma from 3m, 12m and 24m. The vertical axis indicates the level of T (A), DHT (B), and E2 (C). Concentrations of T (A), DHT (B), and E2 (C) in 24m and 12m hippocampus are compared with those in 3m. Data of 3m are taken from Hojo et al., (2009) (Hojo et al 2009). DHT-picolinoyl (m/z = 203), T-picolinoyl (m/z = 253), and E2-PFBz-picolinoyl (m/z = 339) are analyzed by using LC-MS/MS as described in earlier studies (Hojo et al 2009). Data are represented as mean ± SEM. Statistical significance yielded \**p* < 0.05 and \*\**p* < 0.01 vs 3m hippocampus group, and ^##^ *p* < 0.01 vs 3m plasma group.

Importantly, if we use frozen hippocampal tissue, oxidation of 3-OH group of E2 by freeze-thaw processes may cause artificial loss of E2 (Kawato et al 2025). We explain these problems in detail in Supplementary Methods. Therefore, we rapidly extract E2 from freshly separated hippocampal tissue by organic solvent. C18 column and HPLC are also useful for further purification of E2 from contaminating fats. High sensitivity detection was achieved by using picolinoyl derivatization (see Materials and Methods).

Data of steroid concentrations in 3m rats are taken from Hojo et al. (2009) (Hojo et al 2009).

In the hippocampus, the level of T monotonously decreased with age, from 17 ± 2.0 nM (3m) to 5.0 ± 0.9 nM (12m) and 0.2 ± 0.1 nM (24m) (Fig.4). The level of DHT also monotonously decreased with age, from 7.0 ± 2.0 nM (3m) and 2.0 ± 0.4 nM (12m) to 0.4 ± 0.07 nM (24m). The levels of T and DHT in 24m hippocampus extremely decreased to approximately 1% and 4% of those in 3m hippocampus.

The level of hippocampal E2 gradually decreased with age from 8.0± 1.0 (3m) to 8.0± 2.0 (12m) and 2.0 ± 0.5 nM (24m). The E2 level in 24m hippocampus was approx. 25% of that in 3m hippocampus.

Interestingly, the levels of T, DHT and E2 in 3m, 12m and 24 hippocampi were always higher than those in the plasma (Fig. 4). In the plasma, T level decreased from 11 ± 2.0 nM (3m) to 3.0 ± 0.7 nM (12m) and 0.3 ± 0.06 nM (24m). DHT level decreased with age, from 0.6 ± 0.09 nM (3m) to 0.1 ± 0.02 nM (12m) and 0.04 ± 0.004 nM (24m). The levels of T and DHT in 24m plasma were considerably low with approx. 3% and 7%, respectively, of those in 3m plasma. Male plasma E2 levels (0.01 nM at 3m, 0.02 nM at 12m and 0.03 nM at 24m) were always very low and did not show a significant change with aging.

### 2.3. Age-induced change in expression levels of mRNAs encoding sex-steroidogenic enzymes and sex steroid receptors

We performed the PCR analysis in the hippocampus (Hi) for male rats, which are aged 3 month (3m), 12 month (12m), and 24 month (24m). In the following the *italic* name (e.g., *Gapdh*) means the gene name of protein (e.g., GAPDH). The expression level of *Gapdh* mRNA did not show a significant age-related change over 3m, 12, and 24m (Fig. 5A). Therefore, in this section, all the expression levels of enzymes and receptors were normalized by the *Gapdh* level of 3m hippocampus, which was set to be 100%.

**Figure 5:**
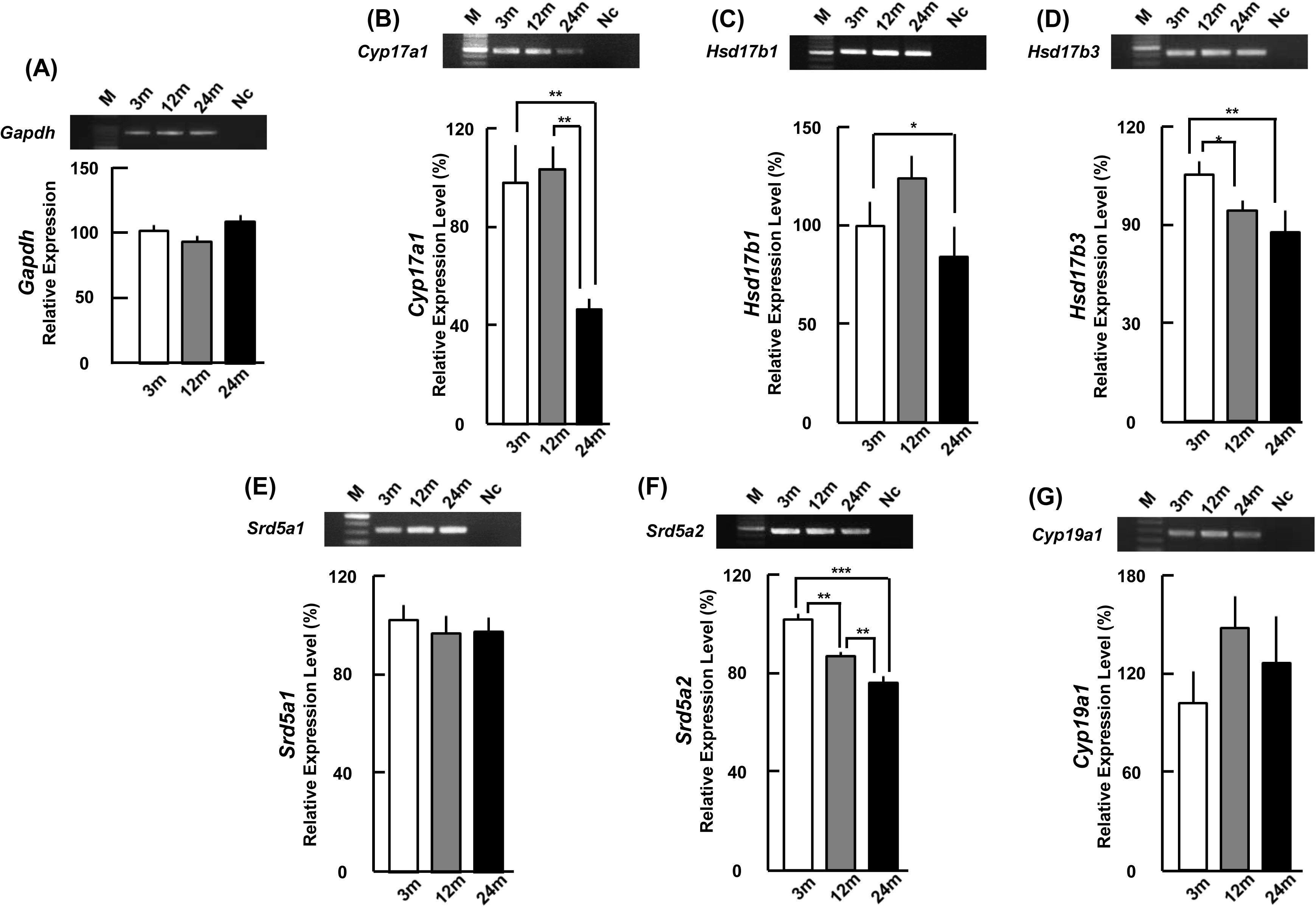
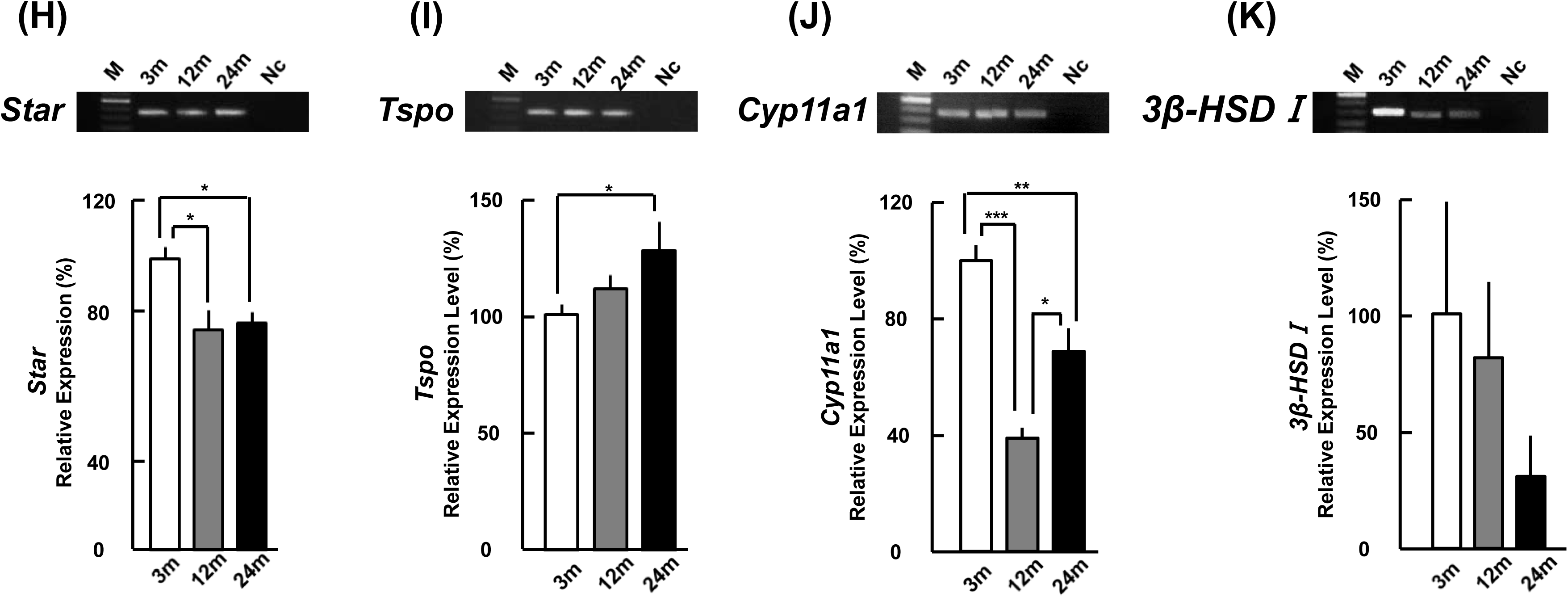
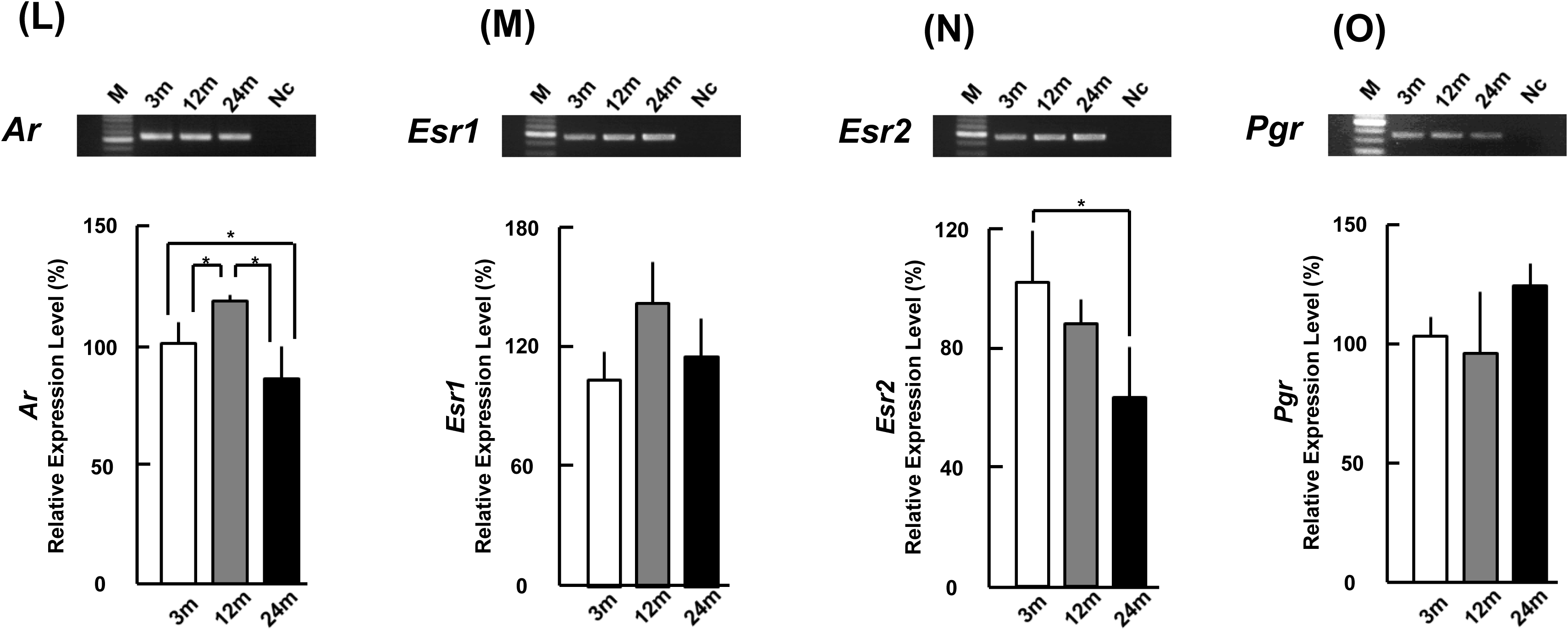
Age-induced change of steroidogenic enzymes, cholesterol transport proteins and steroid receptors in the hippocampus. Upper panels show representative PCR images visualized with ethidium bromide (EtBr) staining. From left to right, 100 bp DNA ladder (M), 3m, 12m, 24m, and the sample without template cDNA as negative control (Nc). Lower panels show mRNA expression levels of the (A) *Gapdh*, (B-K) steroidogenic enzymes ((B) *Cyp17a1*, (C) *Hsd17b1*, (D) *Hsd17b3*, (E) *Srd5a1*, (F) *Srd5a2*, (G) *Cyp19a1*, (H) *Star*, (I) *Tspo*, (J) *Cyp11a1*, (K) *3β-HSDI*), and (L-O) steroid receptors ((L) *Ar*, (M) *Esr1*,(N) *Esr2* and (O) *Pgr*) in 3m, 12m and 24m rats. The vertical axis indicates the relative expression level for each mRNA calculated from the intensity of EtBr bands. (A) The *Gapdh* mRNA expression level was normalized with 3m which is set to be 100%. (B-O) The mRNAs expression levels of each steroidogenic enzymes (B-G, J and K), cholesterol transport proteins (H and I) and steroid receptors (L-O) were normalized with *Gapdh* and the corresponding mRNA expression in 3m rats, as described in Methods. Each value is mean ± SEM. Statistical significance yielded \**p* < 0.05, \*\**p* < 0.01, \*\*\**p* < 0.001. Data are taken from duplicate determinations of 4 rats at each age.

Furthermore, the levels of each enzyme and receptor were normalized by its enzyme and receptor levels in 3m hippocampus, which were set to be 100%, respectively.

#### 2.3.1 Androgen synthesis enzymes and androgen receptor showed age-induced decrease (Figs. 5B-5F)

We investigated the expression levels of androgen synthesis enzymes, including DHEA synthase (P450(17α)), testosterone synthase [17β-hydroxysteroid dehydrogenase (17β-HSD) (1 and 3)], and DHT synthase (5α-reductase (1 and 2)). The level of *Cyp17a1* (mRNA of P450(17α)) decreased with age, from 100% (3m) to 62% (24m) (Fig. 5B). The level of *Hsd17b1* (mRNA of 17β-HSD1) decreased with age, from 100% (3m) to 84% (24m) (Fig. 5C). The level of *Hsd17b3* decreased from 100% (3m) to 76% (24m) (Fig. 4D). Although the *Srd5a1* level did not significantly change with age (Fig. 4E), the level of *Srd5a2* (mRNA of 5α-reductase 2) monotonously decreased from 100% (3m) to 85% (12%) to 75% (24m) (Fig. 5F).

#### 2.3.2. Estrogen synthesis enzyme did not change with age (Fig. 5G)

The expression level of *Cyp19a1* (mRNA of P450arom) did not significantly show age-related change (Fig. 4G), although the *Cyp19a1* level in 12m was a little bit higher than those of 3m and 24m without statistical significance.

#### 2.3.3. Cholesterol transport, synthesis of pregnenolone (PREG) and progesterone (PROG), in upstream of androgen and estrogen synthesis (Figs. 5H-5K)

The level of *Star* (mRNA of steroidogenic acute regulatory protein (StAR)) decreased with age, from 100% (3m) to 75%, and 61% (24m) (Fig. 5H). Interestingly, however, the level of *Tspo* (mRNA of translocator protein (TSPO)), monotonously increased with age, from 100% (3m) to 111% (12m), and 127% (24m) (Fig. 5I).

Pregnenolone synthase *Cyp11a1* (mRNA of P450scc) and progesterone synthase *3β-HSDI* (mRNA of 3β-HSD) were investigated. The expression levels of *Cyp11a1* and *3β-HSDI* were much lower than levels of other sex steroid synthesis enzymes in downstream. The *Cyp11a1* level decreased from 3m (100%) to 69% (24m) in average (Fig. 5J). The age-dependent progressive decrease was observed for *3β-HSDI* level in average (Fig. 5K). In individual experiments, mRNA was not observed (even with 42 PCR cycles) in 2 rats (12m) and 3 rats (24m), and observed in 2 rats (12m) and 1 rat (24m) only, among 4 rats examined.

#### 2.3.4 Age dependence of androgen and estrogen receptors (Figs. 5L-5O)

The expression level of androgen receptor, *Ar*, showed age-induced decrease from 100% (3m) to 85% (24m) (Fig. 5L). Although the level of *Esr1* (mRNA of ERα) did not show age-induced change, the level of *Esr2* (mRNA of ERβ) monotonously decreased from 100% (3m) to 87% (12m) to 62% (24m) (Figs. 5M, 5N). Interestingly, the *Pgr* level (mRNA of progesterone receptor (PR)) did not change with age (Fig. 5O).

#### 2.3.5. Relative comparison of all the different enzymes and receptors at the same age of hippocampus (Fig. 6)

Fig. 6 shows comparison of relative expression levels of all the steroidogenic enzymes and steroid receptors in the same age (24m, 12m and 3m), using equations explained in Methods section. We normalized by setting the level of *Cyp19a1* in 3m hippocampus to be 100%, because the expression level of *Cyp19a1* did not significantly change with age.

**Figure 6:**
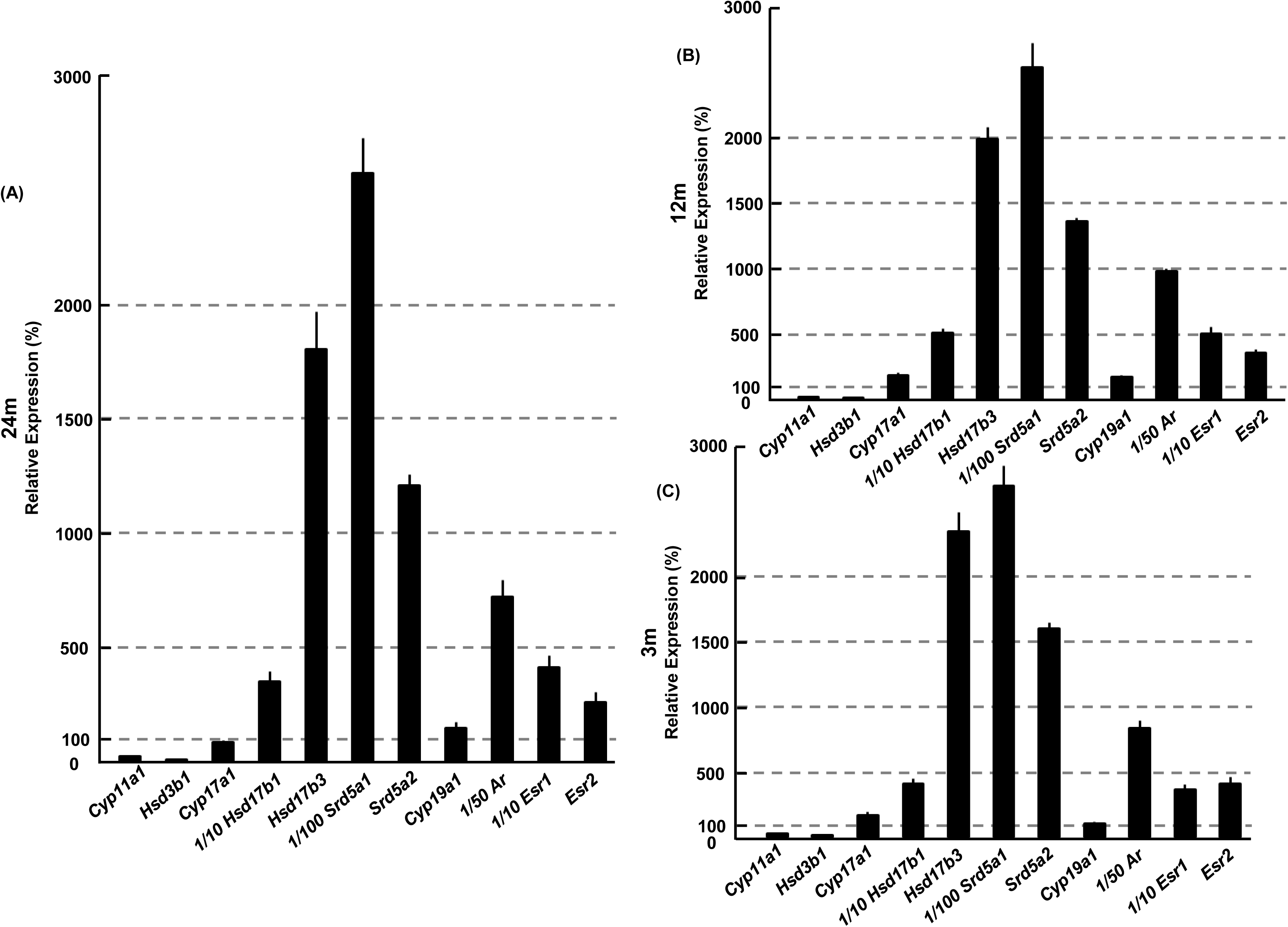
Comprehensive comparison of expression levels of mRNAs between different enzymes and receptors in (A) 24m, (B) 12m and (C) 3m brains. The comparison was performed using the method as described in Materials and Methods. The normalization of each mRNA was performed by setting *Cyp19a1* of 3m Hi as 100%. Data are also normalized by *Gapdh*. The vertical scale is different for each enzymes/receptors. For comparison, the vertical scale should amplify by100-fold for *Srd5a1*; by 50-fold for *Ar*; by10-fold for *Hsd17b1* and *Esr1*; by1-fold for *Cyp17a1*, *Hsd17b3*, *Srd5a2*, *Cyp19a1* and *Esr2*. The data of 3m was modified from Kimoto *et al*. (Kimoto et al 2010).

In order to have clear comparison of many enzymes and receptors whose expression levels are very different, 1/100 fold of *Srd5a1,* 1/50 fold of *Ar,* 1/10 fold of *Hsd17b1* are used for illustration in Fig. 6. Several interesting characteristics were observed at 24m. The expression level of *Srd5a1* was highest among all the enzymes examined. Second highest expression was the *Hsd17b3* level. The expression levels of *Ar* and *Esr1* were relatively high as compared with enzymes. The *Esr1* level was higher (by approxi.15 fold) than the *Esr2* level. The expression levels of *Cyp11a1* and *Hsd3b1* were extremely low among all the enzymes and receptors.

#### 2.3.6. Neurotrophic factors *Bdnf* and senescence marker gene *Sirt1* and *Tp53* (Fig. S2)

*Bdnf* (mRNA of Brain-derived neurotrophic factor (BDNF) and *Brain Insulin-1* (mRNA of Brain Insulin-1) are typical neurotrophic factors in the hippocampus. The *Bdnf* expression level did not show age-related change. The *Brain Insulin-1* level increased with age, from 100% (3m) to 211% (24m). The expression level of *Brain Insulin-1* was much lower (approx. 1/8000) than that of *Bdnf* as judged from different amplification cycles (see Methods).

The *Sirt1* (mRNA of SIRT1, silent information regulator 1) and *Tp53* (mRNA of tumor protein 53) are senescence marker genes. The *Sirt1* level monotonously decreased from 100% (3m) to 80% (12m), and 61% (24m), however, the *Tp53* level did not show age-related change.

#### 2.3.7. Immunostaining of hippocampal enzymes and receptors (Fig. S3)

The localization and expression levels of sex-steroidogenic enzymes and sex-steroid receptors in the hippocampus was investigated in 24m rats by using immunostaining with antibodies (Fig. S3). The characteristic image parameters were compared with those in 3m rats published in our earlier works (Kimoto et al 2001) (Kawato et al 2002) (Hojo et al 2004). Detailed explanation is described in **5.2 Supplementary Results** (Section: Immunostaining of hippocampal enzymes and receptors).

## 3. Discussion

In the field of cognitive decline along with aging, a great volume of data have been accumulated, including aging-induced changes in neurons, synapses and neurotropic factors of the hippocampus and the prefrontal cortex in rodents and nonhuman primates (Buchman et al 2016) (Burke & Barnes 2006) (Hao et al 2007) (Hara et al 2015) (Morrison & Baxter 2012). Particularly, aging dependent dendritic spine loss in the hippocampal pyramidal neurons has been well examined (Dickstein et al 2013) (Hao et al 2006) (Morrison & Baxter 2012). However, neuronal alterations in relation to brain sex steroids not only with normal/healthy aging but also with pathological aging such as AD have not been sufficiently uncovered (Morrison & Baxter 2012) (Carroll et al 2010) (Rosario et al 2010). Therefore, the current observations give a significant contribution to understand what is going on along with healthy/normal aging of the hippocampus, through demonstration of the age-related correlations between the dendritic spine loss and sex-steroids properties (steroid levels, levels of mRNA and protein expression of sex steroidogenic enzymes/receptors). We particularly focused on local hippocampal sex steroids that are candidates for neuroprotective factors in aging and neurodegeneration, because changes in hippocampal sex steroid levels and their functions along with aging have not been extensively investigated.

### 3.1. Decrease in spine density and hippocampal sex steroids with normal aging

The current investigations have demonstrated that age-related degeneration in dendritic spines of male rat hippocampus have close relationships with the decline in androgen action and androgen levels. Along with aging from 3m to 24m, approx. 26% decrease in the total spine density was observed (Fig. 2). With aging, the density of all three types of spines was decreased, and the large-head spine population was most significantly decreased.

Androgen replacement was effective for recovery of the spine density. Sixteen hours after single DHT and T injection, individually, spine density was recovered by approx. 5∼7 % via AR dependent mechanisms in 24m rat hippocampus (Fig. 3). These androgen replacements were significantly effective, because basal hippocampal levels of T and DHT was considerably decreased by going from 3m to 24m as observed by mass-spectroscopic analysis (Fig. 4). Surprisingly, the levels of DHT and T in 24m hippocampus was only approx. 4% and 1%, respectively, of those in 3m hippocampus. These low androgen cases could be assigned to a kind of “andropose state”. This andropose-like state may appear due to not only considerable decrease of testis-derived androgen but also particularly low DHEA production in rat adrenal cortex (containing very low level of cytochrome P450(17α)) which is entirely different from human adrenal cortex that could produce significant level of DHEA even at aged state.

Why the total spine density in 24m hippocampus was maintained at the moderately high density (approx.74% of that of 3m), even with no androgenOne reason may be that the E2 level in 24m hippocampus was still moderately high, approxi.1/4 level of that in 3m, which is effective to keep the moderately high spine density (Hasegawa et al 2015) (Mukai et al 2010). Another reason could be that neurotrophic factors, including brain-derived neurotrophic factor (BDNF), nerve growth factor (NGF) (Perovic et al 2013) and Brain-insulin, were not at all decreased with aging in the hippocampus. These neurotrophic factors are important candidates which could increase spines (Pozzo-Miller et al 1999) and elevate LTP (Kramar et al 2004), but no age-induced decrease in their levels imply no contribution to the observed spine density decline.

It is well known that in AD model mice such as 3xTg-AD and Tg2576 mice, the spine density decreased considerably with aging. These spine degenerations are explained through the effects of Aβ oligomers and Aβ plaque (Bittner et al 2010) (Jacobsen et al 2006), therefore. the cause of spine decrease is very different from healthy aging.

### 3.2. Hippocampal synthesis capacity of androgen and estrogen in aged rats

Until the current investigations, aging-dependence of local androgen and estrogen synthesis systems after adult stages were poorly understood in not only the hippocampus but also other brain regions. Our earlier study (Munetomo et al 2015) described aging dependence of mRNAs of representative steroidogenic P450s and hydroxysteroid-dehydrogenase (HSD) in the hypothalamus (Hy), the Cerebral Cortex (CC) and the Cerebellum (CL). StAR immunostaining had been exceptionally shown in aged rat hippocampal neurons at 24m (Sierra et al 2003).

Local androgen synthesis capacity from PROG to T and DHT depending on P450(17α) and 17β-HSD (1,3) was significantly decreased with aging (Figs. 4, 5), however their mRNA expression levels of *Cyp17a1* and *Hsd17b1,3* in 24m hippocampus still keep approx. 50% − 75% of those in 3m hippocampus. Therefore, local androgen synthesis activity from PROG to T through androstenedione in 24m hippocampus might not be lower than 50% of that in 3m hippocampus (see synthesis pathway in Fig. 1).

However, only a very low level of T (0.2 nM) was observed in 24 hippocampus. Under the assumption that local T synthesis capacity from PROG (see Fig. 1) may be maintained at around 50% as deduced from mRNA levels, why hippocampal T level was extremely low in 24mOne reason of age-dependent considerable decrease in hippocampal T and DHT levels (Fig. 4) is probably a considerable decrease in penetration of circulating T to the hippocampus (down to approx. 3%, by going from 3m to 24m), because circulating T penetrates/permeates into the hippocampus and circulating T accounts for approx. 70 - 80% of hippocampal T, as observed in Fig. 3A and our earlier studies (Hojo et al 2009). It should be noted that circulating DHT level is much lower than hippocampal DHT level (less than 10% of hippocampal DHT level) (Fig. 4), and therefore circulating DHT cannot contribute to maintain hippocampal DHT level. The hippocampal DHT level was maintained by local conversion from hippocampal T (Hojo et al 2009). Another reason of low hippocampal T could be explained as a very efficient local conversion of T to DHT (by 5α-reductase) and T to E2 (by P450arom). T to DHT conversion may be very efficient, because the expression level of 5α-reductase in the hippocampus was the highest among all steroidogenic enzymes (Figs. 4 and 5) and no age dependent decrease was observed for 5α-reductase. Indeed, these efficient rapid conversion from T to DHT and 3a-androstanediol were observed at 3m hippocampus (Hojo et al 2009) (Hojo et al 2011).

T to E2 conversion may also be efficient, because the mRNA level of P450arom did not decrease with aging. Note that no ^3^H-E2 degradation was observed within the hippocampal slices, implying that E2 is very stably present once synthesized (Hojo et al 2004).

Interestingly, E2 synthesis capacity was only moderately decreased even at 24m. *Cyp19* and P450arom immunostaining did not decrease, and therefore the E2 level in 24m hippocampus was still kept ∼1/4 of that in 3m. Penetration of circulating E2 to the hippocampus cannot contribute, because plasma E2 level is always lower than hippocampus E2 level.

The above considerations are based on the sufficient PROG supply to the hippocampus via blood PROG circulation at 24m (Valle et al 2008), and this is supported by the results that plasma PROG level was increase from 6m to 24m in male rats (Valle et al 2008). This is an important point, because local PROG synthesis capacity from cholesterol may be very poor in 24m hippocampus, because expression levels of P450scc and 3β-HSD1 were very low as judged from mRNAs and immunostaining (Fig. 5 and Supplementary Fig. S3).

P450scc mRNA expression level was much lower than mRNAs of other androgen and estrogen synthases. P450scc protein staining was considerably decreased at 24m. 3β-HSD1 mRNA was nearly negligible at 24m. In 3m hippocampus, mRNA expression of 3β-HSD1 was observed for all the 10 rats examined. However, in 24m rats 3β-HSD1 mRNA was not observed in 3 rats (24m), among 4 rats examined. Note that only averaged data are shown in Fig. 4 for 24m. These results imply the almost negligible conversion from PREG to PROG (and from DHEA to T) via 3β-HSD1. From these considerations, aged rat hippocampus needs circulating PROG as an initial precursor steroid for T synthesis, and this is supported by the sufficient level of circulating PROG at 24m (Valle et al 2008).

### 3.3. Comparison between androgen and estrogen receptors with healthy aging of hippocampus

AR expression level was decreased with aging, however the mRNA expression level in 24m hippocampus still keep approx. 85% of that in 3m hippocampus (Fig. 5). On the other hand, T and DHT levels were extremely low in the 24m hippocampus. Therefore, T and DHT supplementation could significantly recover the spine density via AR (Fig. 2). No significant decline was observed in the levels of estrogen receptor ERα with aging (Fig. 5), in contrast to the decrease of AR. Although ERβ mRNA decreased with aging (Fig. 5) (Yamaguchi & Yuri 2012), this ERβ decrease might not significantly alter the synaptic plasticity of the hippocampus. One reason is that the ERβ expression level is only around 10% of ERα expression level (Fig. 5). Another reason is that much smaller contributions of ERβ than those of ERα to spinogenesis and LTP were observed in earlier studies (Hasegawa et al 2015) (Phan et al 2011) (Phan et al 2015).

ERα agonist PPT upregulated spinogenesis and LTP-induction upon weak theta-burst stimulation but ERβ agonist DPN did not show any effects.

Together with no decrease in estrogen synthesis enzyme P450arom, these results suggest that estrogen signaling could still well regulate synaptic plasticity even in aged 24m hippocampus.

### 3.4. No neuron loss but synaptic protein decrease in normal aging

The number of neurons in the hippocampus does not change in healthy aging of rodents, even for rats with impairment of special learning (Rapp & Gallagher 1996) (West et al 1994, West et al 2004). Also no neuron loss occurred in CC via healthy aging (Burke & Barnes 2006) (Kelly et al 2006) (Morrison & Hof 1997) (Morrison & Hof 2002). The number of neurons did not significantly decrease in prefrontal cortex (Peters et al 1994, Smith et al 2004) (Peters et al 1998), visual cortex (Kim et al 1997, Vincent et al 1989), and entorhinal cortex (Gazzaley et al 1997, Merrill et al 2001) of rodents and primates. Therefore, the cognitive deficits, which occurred in healthy aged animals, might be due to decrease in synapses, but not loss of number of neurons. No change in the number of neurons also occurs in Hy during normal aging (Roberts et al 2012). On the other hand, the number of Purkinje cells in CL decreases due to aging (Janmaat et al 2011, Woodruff-Pak et al 2010).

Examination of possible age-induced decrease in hippocampal synaptic proteins is essential for understanding of impairment of cognition. NMDA glutamate receptors (Avila et al 2017, Magnusson et al 2010) and postsynaptic density protein 95 (PSD-95) (Fig. S3) (VanGuilder et al 2011) showed considerable decrease with age, which is a strong evidence for the decline in synaptic plasticity such as LTP with aging.

### 3.5. No reduction in typical endogenous neurotrophic factors

Neurotrophic factors play important roles in synaptic plasticity, maintenance of neural activity and neuroprotection. Importantly in the hippocampus, no age dependent changes in representative neurotrophic factors such as BDNF and NGF have been indicated (Fig. S2) (Croll et al 1998) (Perovic et al 2013).

In Hy and CL, BDNF protein and mRNA level also did not significantly change with age (Munetomo et al 2015) (Ewa et al 2012, Katoh-Semba et al 1998, Perovic et al 2013, Silhol et al 2005).

Brain-Insulin is an another neurotrophic factor (Gerozissis 2003) (de la Monte & Wands 2005). Brain-Insulin increased by going from 3m to 24m in the hippocampus (Fig. S2). Brain-Insulin showed the similar age-dependent increase also in other brain regions including CC, Hy and CL.

These results imply that the currently observed age-dependent spine decrease in the hippocampus cannot be caused by these typical neurotrophic factors, including BDNF and NGF. In AD, although BDNF decreased (Buchman et al 2016), NGF increased (Scott et al 1995). Therefore, age dependent changes in brain neurotrophic factors should be carefully considered.

### 3.6. Age-dependent changes of AR, ER and steroidogenic enzymes in non-hippocampal brain regions

Age-dependent changes of steroid receptors and steroidogenic enzymes showed very different tendency in different brain regions, including the hippocampus, Hy, CC and CL in rats. We discuss typical differences between hippocampus and other regions (Hy, CC, CL) in details in **5.3 Supplementary Discussion**.

### 3.7. Characteristics of hippocampal synaptic degeneration in AD model animals, different from normal aging

Changes in synaptic plasticity and alterations of hippocampal neurons have been extensively investigated in AD model mice as a typical model of pathological aging.

Very different from healthy aging, amyloid β (Aβ) plaques and hyperphosphorylated tau are main factors that affect synapses in AD model mice including triple transgenic AD mice (3xTg-AD) which progressively develop both Aβ and tau pathology in the hippocampus (Bittner et al 2010). In the hippocampus of 3xTg-AD male mice, dendritic spine density declines at 15 months (15m) at which Aβ plaques are abundant (Bittner et al 2010).

Sex-steroids may play a role in neuroprotection in 3xTg-AD mice (Carroll et al 2010) (Rosario et al 2010). In 3xTg-AD male mice, a significant increase in Aβ accumulation in hippocampus was observed in gonadectomized (GDX) mice. Treatment of GDX mice with T or DHT (for 4 m, from 3 m of age until 7 months of age) prevented the increased Aβ accumulation in the hippocampus. E2 also prevented Aβ accumulation in the hippocampus. Levels of tau hyperphosphorylation slightly increased by GDX. Treatment of GDX mice with T and E2 but not DHT reduced tau hyperphosphorylation. These data suggest that T can suppress Aβ pathology through androgen and estrogen pathways and can reduce tau pathology largely through estrogen pathways.

Another type of AD model mice is Tg2576 mice which are human amyloid precursor protein (APP) transgenic mice lines. In our preliminary investigations, a significant spine loss of the total spine density was observed in CA1 region of the hippocampus of 12m Tg2576 female mice, using the 3-dimentional confocal imaging of Lucifer yellow injected hippocampal neurons. From morphology analysis, large-head spines were particularly decreased without significant loss of middle- and small-head spines. The loss of the total spine density in Tg2576 female mice was also observed at 6m and 12m using Golgi stained 2-dimentional analysis (Ricobaraza et al 2012). In Tg2576 male mice, spine loss in dentate gyrus (DG) region was observed at 4m and 12 m using Golgi stained 2-dimentional analysis, however a significant Aβ increase was observed from 18 m (Jacobsen et al 2006). In Tg2576 male, both long-term potentiation (LTP) deficits in hippocampal CA1 and DG and spatial memory deficits in a modified water maze were observed at 6 m. Although Aβ plaques did not develop until 18 m (Hsiao et al 1996), both LTP deficits in hippocampal CA1 and DG and spatial memory deficits in a modified water maze were detected at 6 m. Taken together, these results imply that impaired synaptic plasticity appears before Aβ plaque formation (Chapman et al 1999), therefore normally aged states of 24m hippocampus with spine loss might be similar to pre-Aβ plaque states of AD model animals.

### 3.8. Comparison between semi-quantitative PCR and real-time PCR

We can obtain essentially the same information from the current semi-quantitative PCR method (Hojo et al 2004) (Kimoto et al 2010) and real-time PCR (Munetsuna et al 2009). In order to perform normalization, both methods need to choose a good standard house-keeping gene, which must not change with age in addition to region-independent expression. The real-time PCR also cannot avoid these processes. We observed that *Gapdh* satisfied these criteria. Since the expression levels of mRNAs for steroidogenic enzymes are extremely low in the adult brain (Hojo et al 2004) (Kimoto et al 2010), the primer design is most important. Concerning primer design of steroidogenic enzymes and receptors (with specificity and selectivity), dependent on free energy calculation, our semi-quantitative PCR methods are much more optimized (from many experiences) than real-time PCR (with fewer experiences) (Munetsuna et al 2009). Taken together, as far as we choose the PCR cycle number within the exponential amplification phase, we can achieve essentially the same results between semi-quantitative PCR method and real-time PCR method.

## 4. Materials and Methods

### 4.1. Animals

Male Wistar rats of 3, 12, 24-month-old were used in the current study. Animals were housed in the animal facility of Tokyo Metropolitan Institute of Gerontology, under a 12 h-light / dark cycle and were allowed *ad libitum* access to food and water. All experiments were performed with the approval of the Committee for Animal Research of the University of Tokyo.

### 4.2. Chemicals

DHT and T and Lucifer Yellow were purchased from Sigma-Aldrich (USA). E2, Flutamide and Tamoxifen were from Fujifilm-Wako Chemical (Japan).

Picolinic acid was from Tokyo Chemical Industry (Japan) and [1,2,3,4-^13^C_4_]E_2_ was from Hayashi Pure Chemical (Japan). T-d_3_ and DHT-d_3_ were from CDN Isotope Inc. (Canada). [3H] labeled steroids ([2,4,6,7-^3^H]-E2, [1,2,6,7-^3^H]-T, and [1,2,6,7-^3^H]-DHT) were purchased from Perkin Elmer (USA).

### 4.3. Hippocampal spine analysis

#### Subcutaneous drug administration of animals

DHT (1 mg/kg body weight), T (1 mg/kg body weight), E2 (40 μg/kg body weight), Flu (mg/kg body weight) and Tamoxifen (mg/kg body weight) was dissolved in sesame oil to reach its appropriate concentration. The final volume was adjusted to 400 μl. Subcutaneous administration was performed at 5 pm which was 17 h before the decapitation. During the administration, the rats were gently handled by the experimenter.

#### Slice preparation (from in vivo fixed hippocampus)

Hippocampal slices were prepared from a 12-week-old male rat that was deeply anesthetized and perfused transcardially with PBS (0.1 M phosphate buffer and 0.14 M NaCl, pH 7.3), followed by a fixative solution of 3.5% paraformaldehyde. Immediately after decapitation, the brain was removed from the skull and post-fixed with the fixative solution. The dorsal hippocampus was then dissected and 400 µm thick transverse slices to the long axis, from the middle third of the hippocampus, were prepared with a vibratome (Dosaka, Japan).

#### Imaging and analysis of dendritic spine density and morphology

##### Current injection of Lucifer Yellow

Neurons within slices were visualized by an injection of Lucifer Yellow under a Nikon E600FN microscope (Japan) equipped with a C2400-79H infrared camera (Hamamatsu Photonics, Japan) and 40× water immersion lens (Nikon). A glass electrode was filled with 4% Lucifer Yellow, which was then injected for 5 min using Axopatch 200B (Axon Instruments, USA). With this process, approx. five neurons within a 10–20 μm depth from the surface of a slice were injected.

##### Confocal laser microscopic imaging and analysis

The imaging was performed from sequential z-series scans with LSM5 PASCAL confocal microscope (Zeiss, Germany) at high zoom (3.0) with a 63× water immersion lens, NA 1.2 (Zeiss). For Lucifer Yellow, the excitation and emission wave lengths were 488 nm and 530 nm, respectively. For analysis of spines, three-dimensional image was reconstructed from approximately 30 sequential z-series sections for every 0.45 μm. The applied zoom factor (3.0) yielded 23 pixels per 1 μm. The confocal lateral resolution was approximately 0.16 μm. The z-axis resolution was approximately 0.47 µm. The confocal lateral resolution was approximately 0.16 μm. The z-axis resolution was approximately 0.47 µm. Our resolution limits were regarded to be sufficient to allow the determination of the head diameter of spines in addition to the density of spines. Confocal images were then deconvoluted using AutoDeblur software (AutoQuant, USA).

The density of spines as well as the head diameter were analyzed with Spiso-3D (automated software calculating mathematically geometrical parameters of spines) developed by Bioinformatics Project of Kawato’s group (Mukai et al 2011). Spiso-3D has an equivalent capacity with Neurolucida (MicroBrightField, USA), furthermore, Spiso-3D considerably reduces human errors and experimenter labor. The single apical dendrite was analyzed separately. The spine density was calculated from the number of spines along secondary dendrites having a total length of 40-60 µm. These dendrites were present within the stratum radiatum, between 100-200 µm from the soma.

Spine shapes were classified into three categories as follows: (1) small-head spines, with a head diameter between 0.2-0.4 µm; (2) middle-head spines, which have 0.4-0.5 µm spine heads; and (3) large-head spines, with a head diameter between 0.5 and 1.2 µm. The reason of this classification depends on the consideration that these three types of spines may have different efficiency in signal transduction. Small-, middle-, and large-head spines probably have different number of α-amino-3-hydroxy-5-methyl-4-isoxazolepropionic acid (AMPA) receptors, and therefore these three types of spines might have different efficiency in memory storage. The number of AMPA receptors (including GluR1 subunits) in the spine increases as the size of postsynapse increases, whereas the number of N-methyl-D-aspartate (NMDA) receptors (including NR2B subunits) might be relatively constant (Shinohara et al 2008). Because the majority of spines (> 93 %) had a distinct head, and stubby spines and filopodia did not contribute much to overall spine distribution, we analyzed spines having a distinct head.

### 4.4. Mass-spectrometric assay of steroids

Detailed procedures are described in our previous publications (Hojo et al 2009). Briefly, the analysis consists of three steps described in the following:

#### First Step: Purification of steroids from hippocampi with normal phase HPLC

We performed sequential preparations, including homogenization of freshly isolated hippocampal tissues (not frozen-thawed tissues), organic solvent extraction of steroids, removal of fats with C18 column, and purification of T, DHT, E2 with normal phase HPLC (Hojo et al 2009). Importantly in frozen-thawed tissues, oxidation of 3-OH group of E2 may cause artificial modification of E2, because attack by reactive oxygens species (produced from mitochondria) may occur in freezing and thawing processes because of leaking and losing of antioxidants (such as glutathione and vitamin C) and antioxidant enzymes (such as superoxide dismutase and glutathione peroxidase) due to braking of cell membranes by crystallization of intracellular water and melting of intracellular ice. Therefore, we rapidly extract E2 from freshly separated hippocampal tissue by organic solvent (ethyl acetate/hexane).

#### Second Step: Derivatization of HPLC-purified steroids before application to LC (reverse-phase)-MS/MS

We prepared T-17-picolinoyl-ester, DHT-17-picolinoyl-ester and E2-pentafluorobenzyl (PFBz)-17-picolinoyl-ester by derivatization of T and DHT and E2 (Hojo et al 2009) (Hojo et al 2011). Derivatized T, DHT and E2 with picolinoyl is useful for induced-ionization by electron spray of MS/MS. Derivatization of E2 with pentafluorobenzyl is useful for improving evaporation efficiency by electron spray.

#### Third Step: Determination of the concentration for T, DHT and E2 using LC-MS/MS

For determination of the concentration of T, DHT and E2, the liquid chromatography-tandem mass spectrometry (LC-MS/MS) system, which consists of a Shimadzu LC system and POI-5000 triple stage quadrupole mass spectrometer (Applied Biosystems, USA) were employed. In the multiple reaction monitoring mode, the instrument monitored the m/z transition, from 394 to 253 for T-picolinoyl, from 396 to 203 for DHT-picolinoyl and from 558 to 339 for E2-PFBz-picolinoyl, respectively (see Fig. S1). Isotope-labeled steroid derivatives were used for internal standards in order to calibrate the retention time as well as measure recovery of steroids.

The limits of quantification for steroids were measured with blank samples, prepared alongside hippocampal samples through the whole extraction, fractionation and purification procedures. The limits of quantification for T, DHT, E2 were 1 pg, 1 pg, 0.3 pg per 0.1g of hippocampal tissue or 1 mL of plasma, respectively (Table S1). From the calibration curve using standard steroids dissolved in blank samples, the good linearity was observed.

### 4.5. Molecular biological analysis of steroidogenic enzymes and steroid receptors

#### Total RNA isolation

Rats were deeply anesthetized and were decapitated. The hippocampi were quickly removed and immersed in ice-cold artificial cerebrospinal fluid (pH 7.4, 290 mOsm), and stored in liquid nitrogen until use. Total RNA was extracted using SV Total Isolation System (Promega, USA) according to the manufacturer’s instructions. Total RNA was then treated with Recombinant DNase I (RNase-free DNase I; Takara, Japan), and then purified. The concentration was quantified by absorption at 260 and 280 nm.

#### Primer design (Table S2AB)

Since the expression of steroidogenic enzymes and steroid receptors is extremely low in the brain, PCR primers with high sensitivity and specificity were carefully designed for precise analysis (Kimoto et al 2010). We designed the primers considering Gibbs energy (ΔG) as follows:

1. ΔG of the whole primer (ΔG_(whole)_) was calculated with a nearest-neighbor model, and set to be below the mean value for all of the primer candidates (ΔG_av_) to obtain good stability in primer-target interaction. (2) ΔG of the five 3’-terminal bases of the primer (ΔG_(5base)_) was set to be higher than ΔG_av_ for improved specificity. Consequently, ΔG of 5’-terminal bases was set to be less than the ΔG_av_, resulting not only in the improved stability in 5’-terminal interaction but also in the improved DNA polymerase recognition. Two (forward and reverse) primers were designed on separate exons and primers were designed to avoid ‘PCR debris’ including primer dimers.

#### RT-PCR

Semi-quantitative PCR was performed essentially as described in our earlier study (Hojo et al 2004) (Kimoto et al 2010). As we explain in Discussion 4.9., we can obtain essentially the same information from the current semi-quantitative PCR method and real-time PCR (Munetsuna et al 2009).

The amount of cDNA, the number of PCR cycles, and the sequences of oligonucleotide primers used in RT-PCR are shown in Supplementary Table S2A, S2B.

For RT, total RNA was reverse-transcribed into first-strand cDNA using an oligo(dT) primer. Reaction solution (25 μL) contained 10 μg of total RNA, 1x RT buffer, 1 mM dNTP mixture, 2 μg oligo(dT)_15_ (Promega, USA), 40 U RNasin *Plus* (Promega), and 200 U RTase (Toyobo, Japan). The reaction was carried out at 42°C for 60 min, and stopped by heating 75°C for 15 min. cDNA was treated with 4 U RNase H (Takara bio, Japan) at 37°C for 30 min, and stored at -20°C until use.

PCR was performed in 25 μL of PCR mixture comprising cDNA corresponding to 100 ng of total RNA, 1 x PCR buffer, 0.2 M dNTP mixture, 0.2 μM forward and reverse primers, and 0.63 U Blend Taq polymerase (Toyobo, Japan). PCR was performed with cycle reactions at 95°C for 30 sec, 56-68°C for 20 sec, and 72°C for 30 sec, with an initial denaturing at 95°C for 2 min and a final elongation at 72°C for 5 min. The PCR products were applied to 2% agarose gels. Gels were stained with ethidium bromide (EtBr) and visualized under UV light. Fluorescence images were recorded with Printgraph (ATTO, Japan). For quantitative estimation, images of the bands were analyzed using the Image J software (National Institutes of Health, Bethesda, MD). In all the cases, we first plotted amplification curves in order to obtain the exponential amplification phase of PCR plot.

#### DNA sequencing

PCR products were extracted from agarose gels using a Wizard SV Gel and PCR Clean-up System (Promega) and cloned into pGEM-T-Easy vectors (Promega, USA). Sequencing reactions were performed using a BigDye Terminator v3.1 Cycle Sequencing Kit (Applied Biosystems, Foster City, CA, USA). Signals were detected using an ABI PRISM 3130 Genetic Analyzer (Applied Biosystems, USA). In all the expression analyses, the sequence identity between PCR products and target sequences was confirmed with DNA sequencing.

#### Comparison of the mRNA levels for different enzymes and receptors

We used the comparison method for mRNA levels of different enzymes/receptors obtained by using different primers (Kimoto et al 2010). Relative abundance of different genes was estimated by adopting the expression level of glyceralaldehyde-3-phosphate dehydrogenase mRNA (*Gapdh*) as an internal standard. Optical density value in each band was divided by the number of (1+e)^c^, where c is a PCR cycle number and e is an amplification efficacy obtained from the PCR amplification curves in the exponential amplification phase. More theoretical details were described in (Kimoto et al 2010).

### 4.6. Immunohistochemistory

Immunohistochemical staining was performed essentially as described in our earlier studies (Kimoto et al 2001) (Hojo et al 2004). The hippocampi were frozen-sliced with a cryostat. After application of primary antibody, the slices were incubated for 16-48 h. Primary antibodies used are rabbit anti-P450scc IgG (aa421-441,Chemicon) (1:200), rabbit anti-StAR IgG (PA1-560), Affinity BioReagents) (1:500), rabbit anti-AR IgG (PG-21, Millipore) (1:5000), rabbit anti-ERα IgG (1:1000), rabbit anti-guinea pig P450(17α) IgG (Shinzawa et al 1988) (1:1000), rabbit anti-human P450arom IgG (Jakab et al 1993) (1:1000), and mouse monoclonal anti-PSD95 IgG (ab2723, abcam) (1:1000). For secondary antibody, biotinylated goat anti-rabbit IgG and streptavidin-horseradish peroxidase complex (Vector Lab.) were applied. For mouse monoclonal antibody, biotinylated anti-mouse IgG was used as secondary antibody. Immunoreactive cells were detected with diaminobenzidine-nickel staining.

### 4.7. Statistical analysis of hippocampal dendritic spines and mRNA expression

For dendritic spine analysis, drug-treated dendrite images were used for spine analysis, and typical images were shown in Figs 2 and 3. Each dendrite has approx. 50 μm in length including approx. 75-120 spines. In Figs 2 and 3, for each steroid and inhibitor administration analysis of 24m rat dendritic spines, we used 3 rats, 12 slices, 24 neurons, 48 dendrites, and approx. 4100 spines. For statistical analysis of spines in Fig.2, we employed two-way ANOVA, followed by Tukey-Kramer multiple comparison’s test.

For statistical analysis of age dependent changes of steroid levels in Fig.4 and mRNA expressions in Figs.5 and S2, we employ one-way ANOVA, followed by Tukey-Kramer multiple comparison’s test.

## 5. Supplementary materials

### 5.1 Supplementary methods

#### 5.1.1 Mass-spectrometric assay of steroids in early postnatal and developing animals

Additionally, postnatal 1 (P1), postnatal 10 (P10) and 4 week-old male Wistar rats were also used to determine concentrations of T and E2 in order to compare them with those in 3m, 12m and 24m. Male rats, including pregnant mother rats, were purchased from Tokyo Experimental Animals Supply (Japan). All these animals were maintained under a 12 h light/12 dark exposure and free access to food and water. The experimental procedure of this research was approved by the Committee for Animal Research of Univ of Tokyo. Purification of steroids and LC-MS/MS determination are the same as young and aged hippocampi. Results are shown in Supplementally Table S3.

#### 5.1.2 Difference in E2 concentration between fresh and frozen thawed hippocampus

For assay of hippocampal E2 concentration, freshly prepared hippocampus (but not frozen-thawed hippocampus) must be used (Kawato et al 2025). We here explain the reason. The freeze-thaw processes using deep freezer (-80 degree) may damage brain tissues and cells due to crystallization of intercellular water (in freezing processes) and melting of intercellular ice (in thawing processes). Crystallization of intercellular water by freezing may break cell membranes, leading to leaking and losing of antioxidants (such as glutathione and vitamin C) and antioxidant enzymes (such as superoxide dismutase and glutathione peroxidase). Reactive oxygen species produced from mitochondria may easily attack lipids in cell membranes in the absence of these antioxidants and antioxidant enzymes. Melting of intercellular ices may also break cell membranes, leading to leaking and losing of antioxidants and antioxidant enzymes.

Under these bad conditions, membrane lipid peroxidation would occur, leading to oxidation of OH at C-3 position of E2, resulting in modification of E2 molecules. Resultant E2 concentration may become 0.05 - 0.1 nM in adult male rats (Kawato et al 2025). In this way, the hippocampal E2 concentration may significantly decrease in frozen-thawed hippocampal tissues.

In this way, using freshly prepared (but not frozen-thawed) hippocampal tissues, we can always observe the E2 concentration higher than 3 nM in adult 3m male rats (Hojo et al 2009), (Kato et al 2013).

Interestingly and surprisingly, hippocampal T concentration was not considerably different between frozen-thawed hippocampus and freshly prepared hippocampus (Kawato et al 2025). This is because, T has an oxygen (O) at C-3 position which is resistant to oxidation and inactivation. Therefore, T molecules might not be modified by freeze-thaw processes of hippocampal tissues.

### 5.2 Supplementary Results

#### Immunostaining of hippocampal enzymes and receptors (Fig. S3)

In 24m rats, cytochrome P450(17α) and cytochrome P450arom were expressed in pyramidal neurons of CA1-CA3 and granule cells in dentate gyrus (DG) (Fig. S3). The distributions of these enzyme were similar to those obtained in 3m rats (Kawato et al 2002) (Hojo et al 2004). The stained intensity of P450(17α) and P450arom was not significantly changed between 24m and 3m (Hojo et al 2004). On the other hand, the stained intensity of P450scc at 24m was dramatically weaker than that in 3m (Kimoto et al 2001). The stained intensity of PSD95, synaptic marker protein, very weak at 24m, which is a good agreement with earlier reports (Fig. S3) (VanGuilder et al 2010).

Since immunostaining intensity does not have enough sensitivity, almost no significant changes were observed in the expression levels of sex-steroid receptors (AR and ERα) between 24m and 3m (Fig. S3) (Mukai et al 2007). In 24m hippocampal slices, ERα was expressed with almost the same level in pyramidal neurons of CA1-CA3 regions and granule cell in DG, however, AR expression level was the strongest in CA1, and almost no expression in DG (Hojo et al 2014).

It should be noted that the quantitativity of expression levels from immunostaining analysis is low, therefore, less than ∼30% difference in expression level cannot be evaluated with immunostaining data.

### 5.3 Supplementary Discussion

#### 5.3.1 Age-dependent changes of AR, ER and steroidogenic enzymes in non-hippocampal brain regions

Age-dependent changes of steroid receptors and steroidogenic enzymes showed very different tendency in different brain regions, including the hippocampus, Hy, CC and CL in rats.

Hy has the highest expressions of steroid receptors (AR and ER) and steroidogenic enzymes among other regions (hippocampus, Hy, CC and CL), and often they showed no decline with Hy aging (Munetomo et al 2015).

With aging of Hy, mRNAs of AR and ERα increased, and mRNA of ERβ, P450arom and 5α-reductase did not decrease by going from 3m to 24m (Munetomo et al 2015). In Hy, the number of ERα–immunoreactive cells and AR-immunoreactive cells increased with age (Wu et al 2009).

Since the sealing of blood-brain-barrier in Hy is much looser than that in other brain regions, including hippocampus, CC and CL, the decline in plasma sex steroids may induce resistance against the tendency of age-induced decrease in receptors and steroidogenic enzymes in Hy. These age-resistant phenomena might occur, through hypothalamus-pituitary-gonadal axis during aging processes, due to strong interactions between plasma sex-steroids and Hy.

The expression levels of ERα mRNA increased in CC from 3m to 24m, but did not significantly change in CL as similar to the hippocampus (Munetomo et al 2015).

On the other hand, AR mRNA level decreased in CC at 24m, but did not change in CL with age (Munetomo et al 2015).

With aging, the expression level of ERβ mRNA did not change in CC, but decreased in CL (Munetomo et al 2015). ERβ mRNA-positive cells, however, decreased in CC layer 6 and amygdala (Yamaguchi & Yuri 2012).

Interestingly, in CC and CL, almost no expression of P450arom mRNA was observed over 3m − 24m (Kimoto et al 2010) (Munetomo et al 2015), implying that the activity of estrogen synthesis in CC and CL would be very weak.

CC and CL of young adult and aged rats might need E2 supply from other regions, including Hy or hippocampus, via diffusion/penetration of E2 through cerebrospinal fluid, since hippocampus and Hy could have relatively high activity of E2 synthesis (Hojo et al 2009) (Kimoto et al 2010). Penetration of low level E2 (∼ 0.02 nM) from blood plasma may not be sufficient in the adult and aged male brain. On the other hand, E2 may be sufficiently present in CL in neonatal stage, since *Cyp19a1* is significantly expressed in CL at neonatal stage (Sakamoto et al 2003). Note that the expression level of P450arom mRNA in Hy is approxi. 2-fold of that in the hippocampus and did not decrease with age (Munetomo et al 2015).

#### 5.3.2 Difference in hippocampal T levels between rat and human in old age

The reason why the level of T in the hippocampus and plasma decreased to extremely low level at 24m (Fig. 3) may be due to extremely poor adrenal-synthesized DHEA in rats, because of lacking cytochrome P450(17α) in the adrenal glands. (Le Goascogne et al 1991). Aging induced impairment of DHEA and T synthesis within rat testis directly decreases T levels of the plasma and the hippocampus, because 70 – 80% hippocampal T is derived from the blood circulation (Hojo et al 2009).

On the other hand, human adrenals can produce high level of DHEA because of sufficient expression of adrenal cytochrome P450(17α). This DHEA would be converted to T in other peripheral organs. Therefore, human plasma T did not decrease extremely, but moderately decreased along with aging (Travison et al 2007). Difference in aged T effects on rodents and human should be carefully considered.

---

**Supplementary Figure S1.**
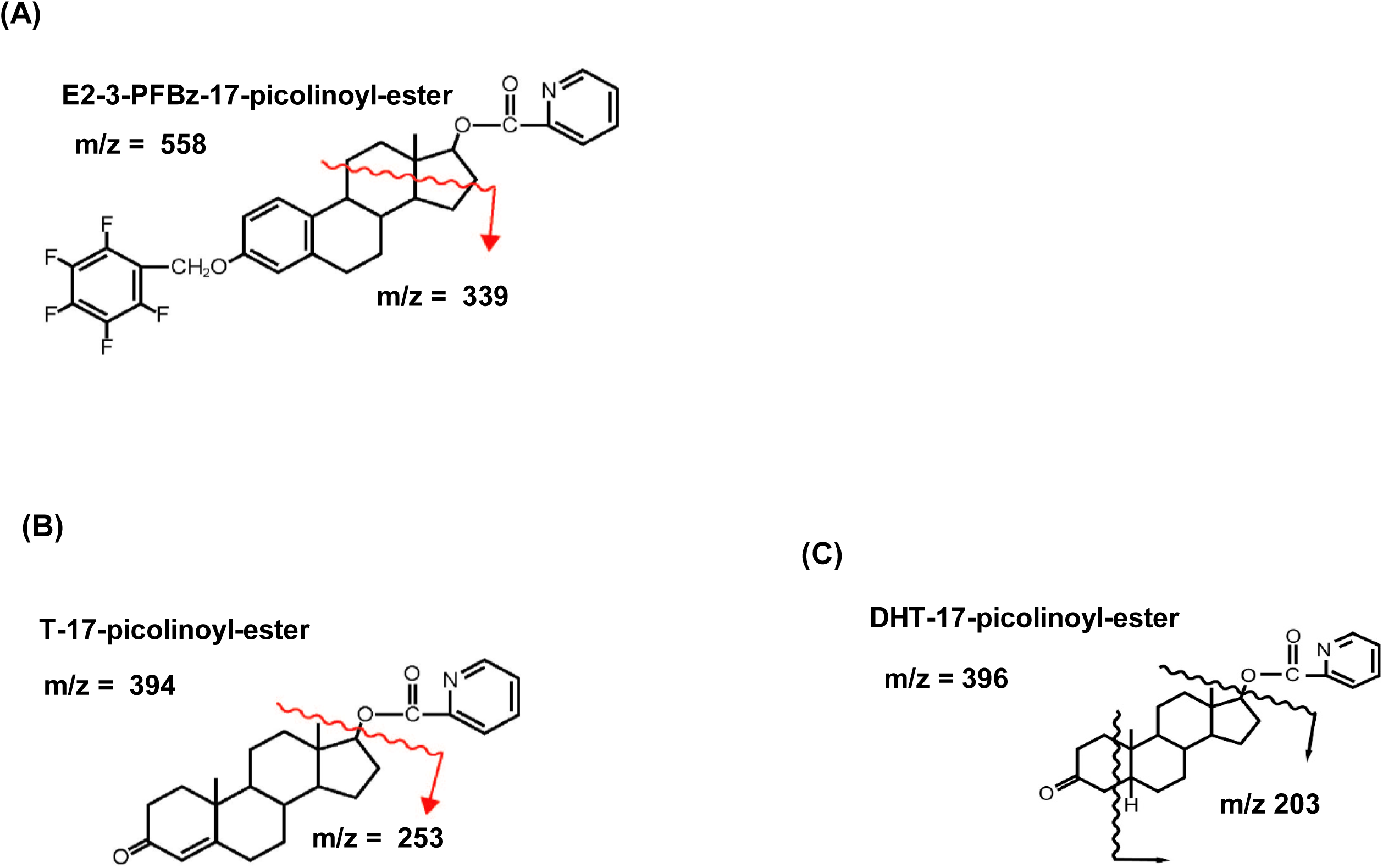
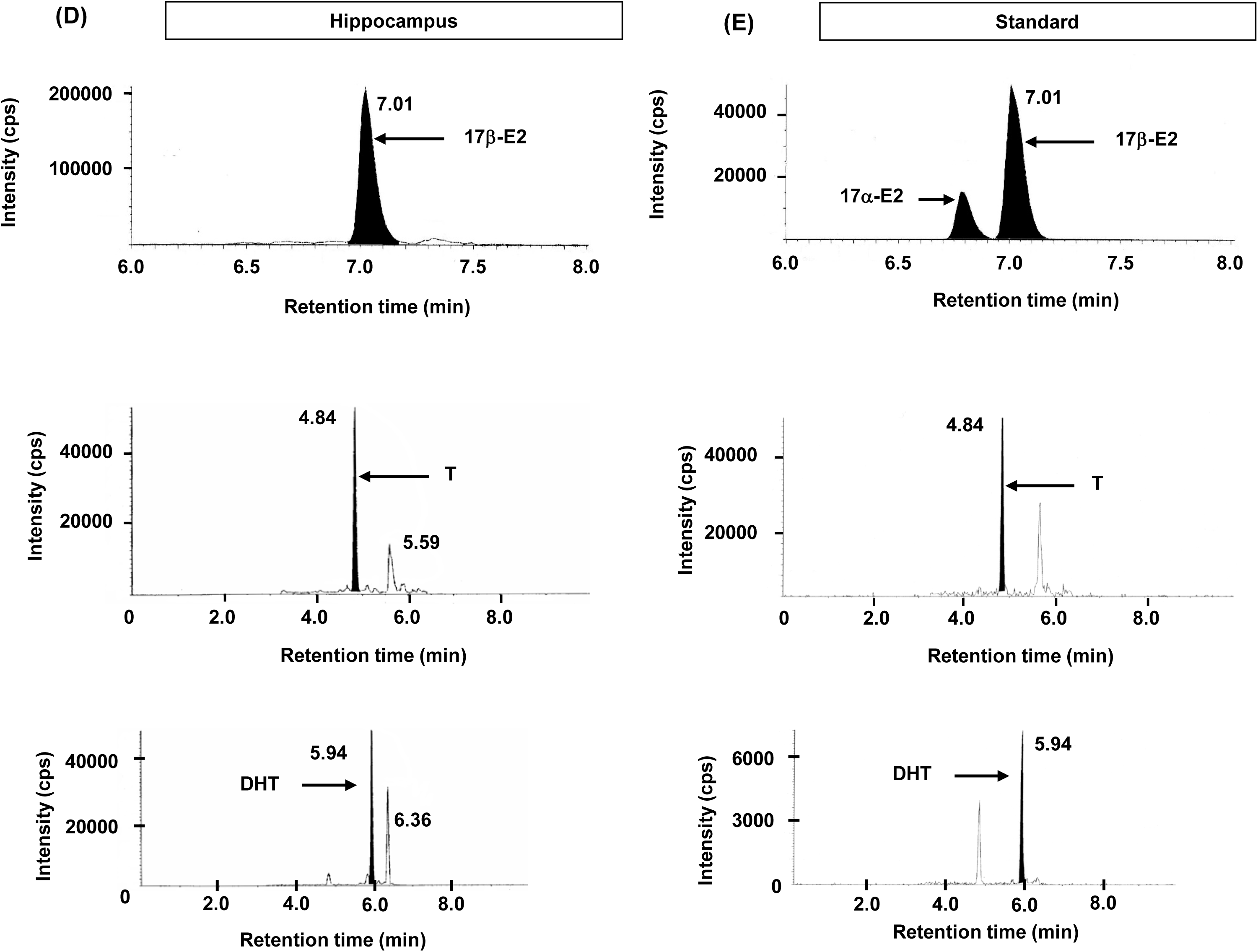
: Steroid derivatives and their fragmented ions used for analysis with LC-MS/MS. (A) E2-3-pentafluorobenzoxy-17-picolinoyl-ester (m/z = 558) and its fragmented ion (m/z = 339), (B) T-17-picolinoyl-ester (m/z = 394) and its fragmented ion (m/z = 253), (C) DHT-17-picolinoyl-ster (m/z = 396) and its fragmented ion (m/z = 203). (D, E) Chromatographic profiles showing the retention time of the fragmented ions of E2-PFBz-picolinoyl, T-picolinoyl and DHT-picolinoyl, with the m/z transition, from 558 to 339 for E2-PFBz-picolinoyl, from 394 to 253 for T-picolinoyl, from 396 to 203 for DHT-picolinoyl, respectively. For these steroids, the retention time of the hippocampal steroid peak (D) was the same as that of standard steroid-picolinoyl (E). (E, F, G) Calibration curves for LC-MS/MS using standard steroids dissolved in ethanol. Horizontal (x) axis indicates the concentration of added standard steroid. Vertical (y) axis indicates the relative intensity obtained from the chromatogram. (A) Calibration curve for E2. Linearity was observed between 0.1 pg/mL to 1000 pg/mL (in this figure only until 400 pg/mL is shown). (B) Calibration curve for T. Linearity was observed between 0.5 pg/mL to 1000 pg/mL. (C) Calibration curve for DHT. Linearity was observed between 0.5 pg/mL to 1000 pg/mL.

**Supplementary Figure S2:**
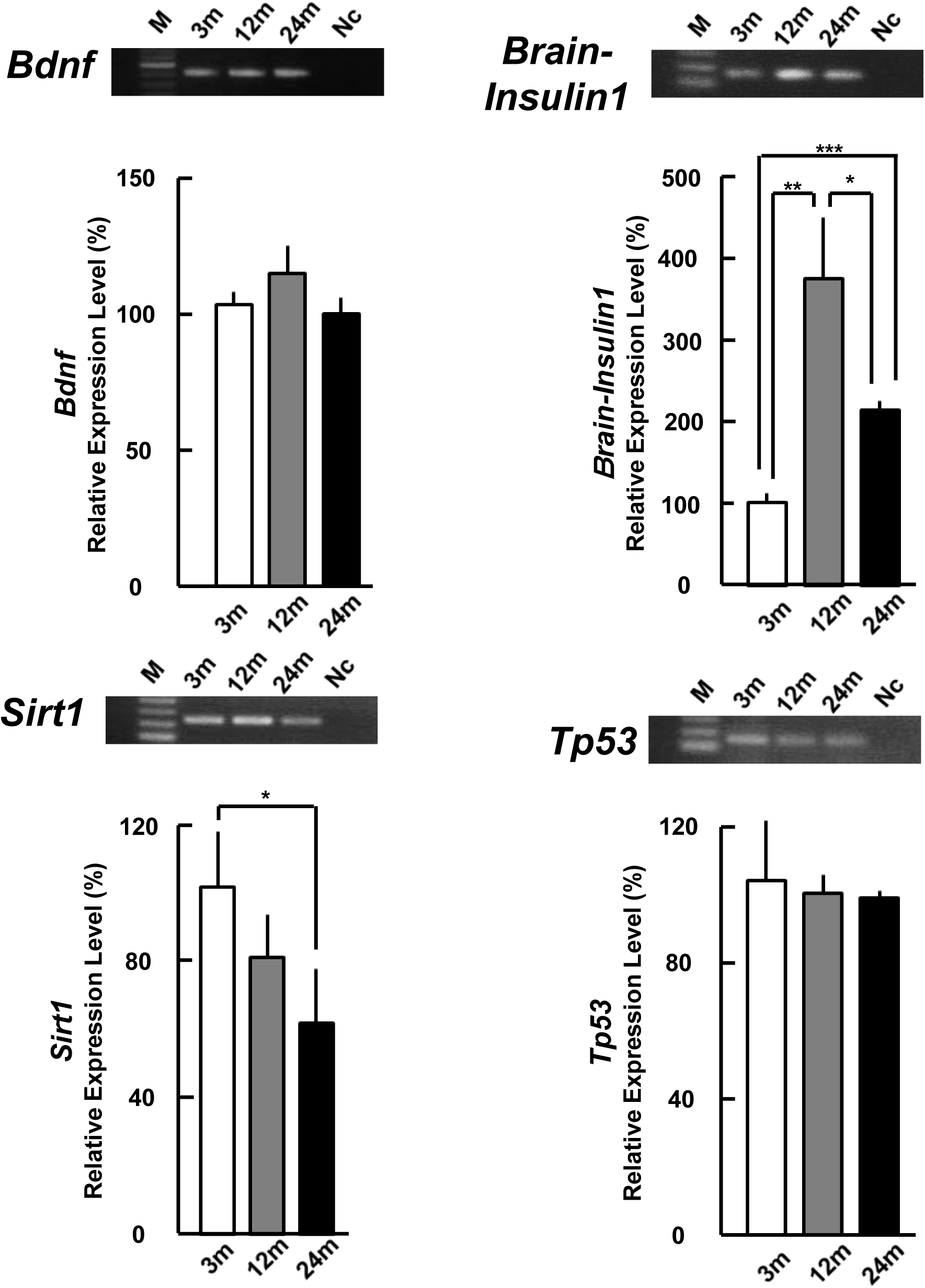
The expression levels of the neurotrophic factors (*Bdnf* and *Brain-Insulin1*), and senescence marker genes (*Sirt1* and *Tp53*) in the hippocampus. Upper panels show typical PCR images. From left to right, 100 bp DNA ladder (M), 3m, 12m, 24m, and the sample without template cDNA as negative control (Nc). Lower panels show mRNA expression levels of (A) *Bdnf* and (B) *Brain-Insulin1*, and (C) *Sirt1* and (D) *Tp53* at 3m, 12m and 24m. The vertical axis indicates the expression level for each mRNAs calculated from the intensity of EtBr bands. Each value is mean ± SEM. Statistical significance, *p < 0.05, **p < 0.01, ***p < 0.001. Data are taken from duplicate determinations for 4 rats at each age. The normalization of each mRNAs was performed in the same manner as Fig. 4.

**Supplementary Figure S3:**
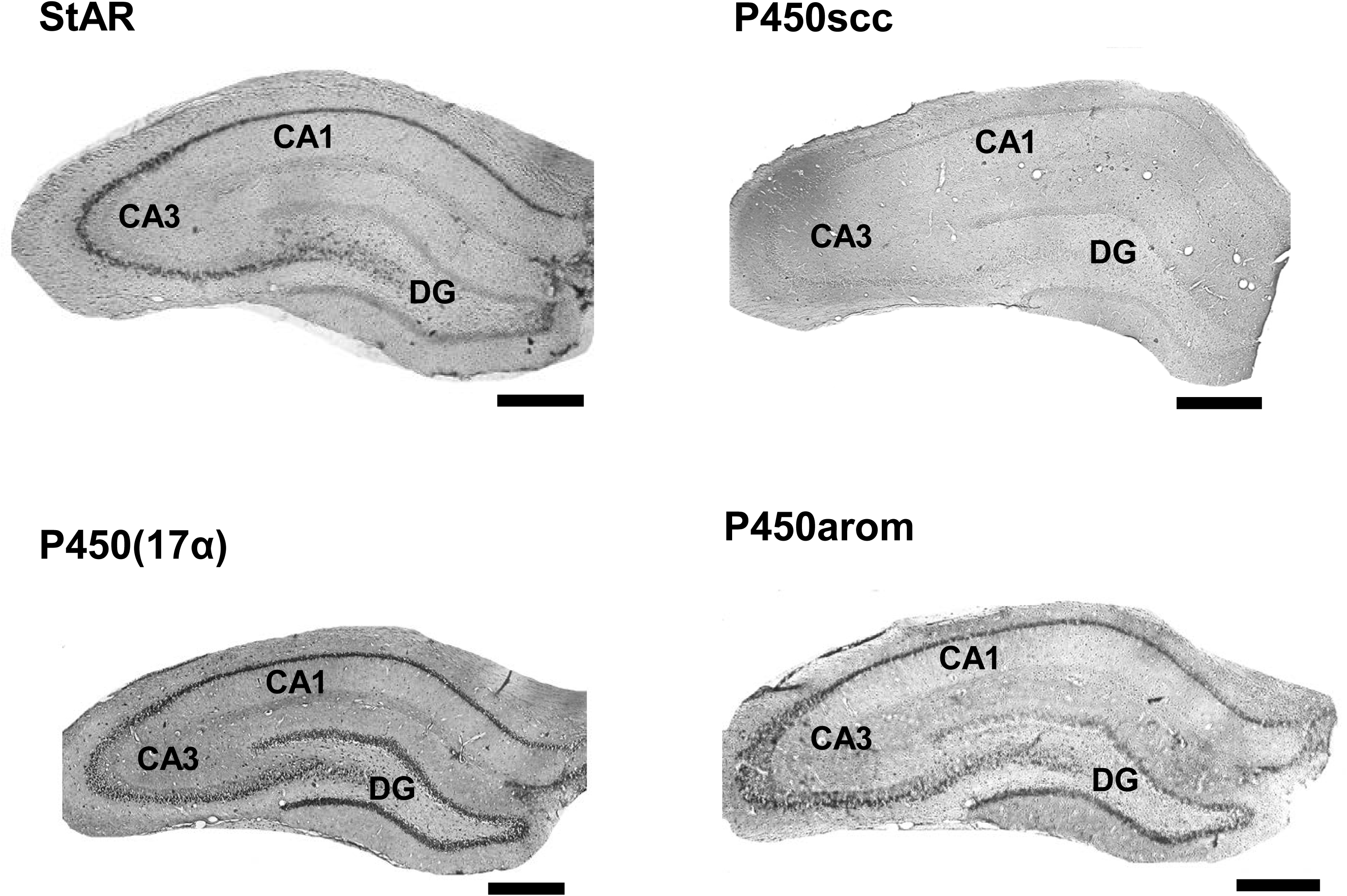
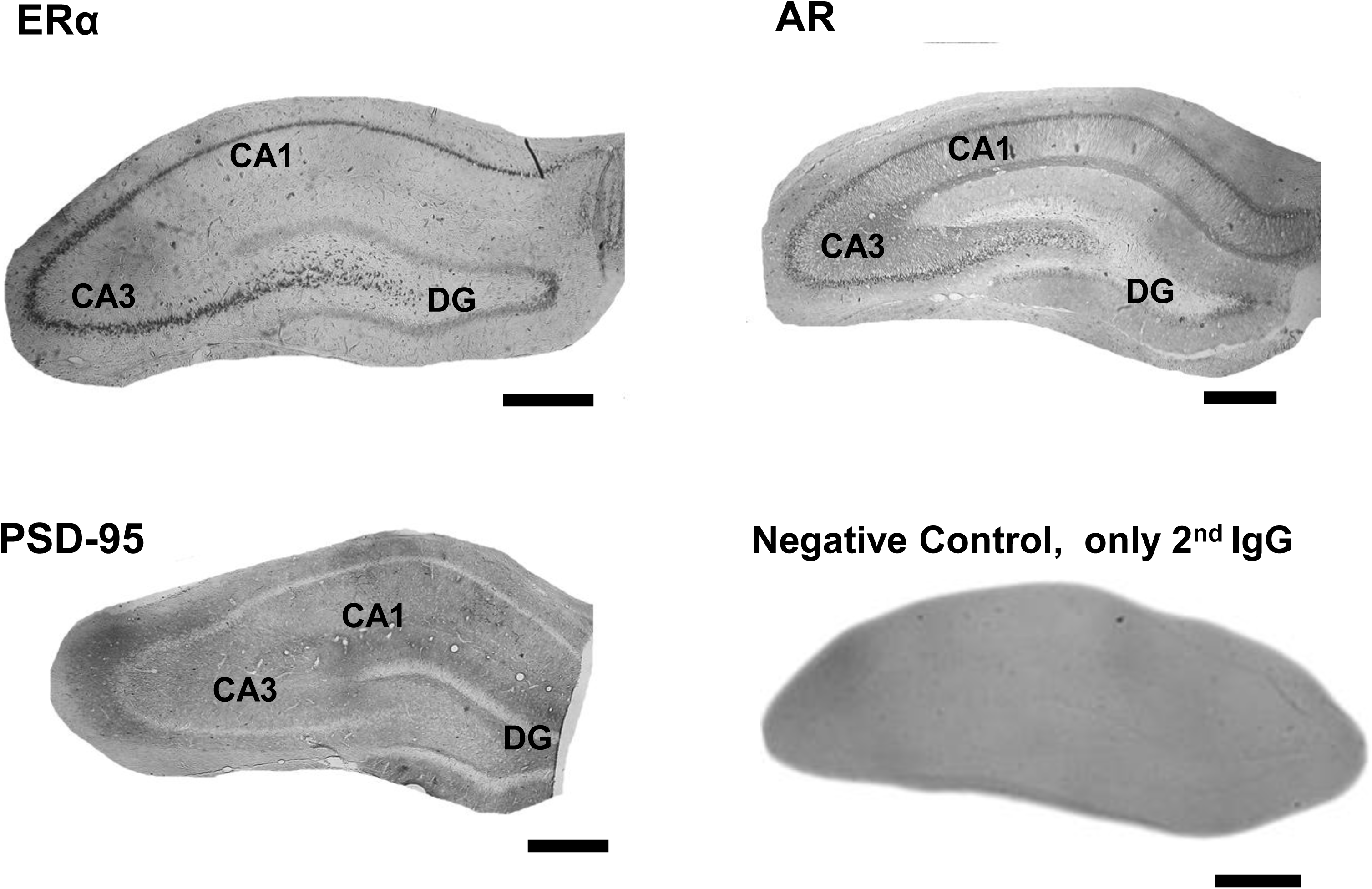
Immunostaining of sex steroidogenic enzymes and receptors in the 24m hippocampal slices. Moderate expressions of StAR, P450(17α), P450arom, ERα, AR are shown in CA1, CA3 and DG regions. AR is mainly expressed in CA1. In contrast, negligible expressions of P450scc and PSD95 are shown. Scale bar 500 μm.

**Table S1.**
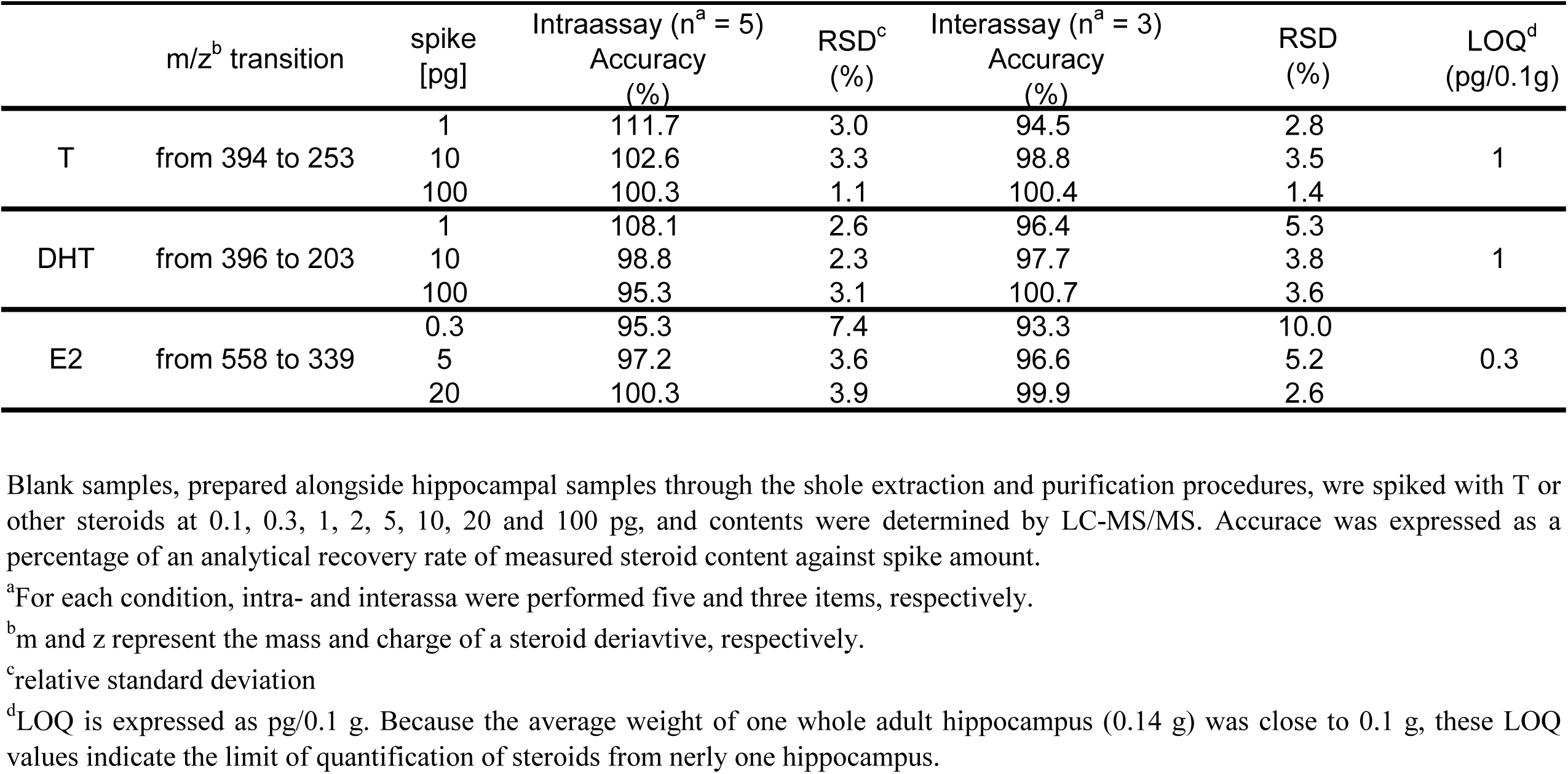
The intra- and inter-assay of accuracy and precision as well as the limit of quantification (LOQ) for each steroid.

**Table S2A.**
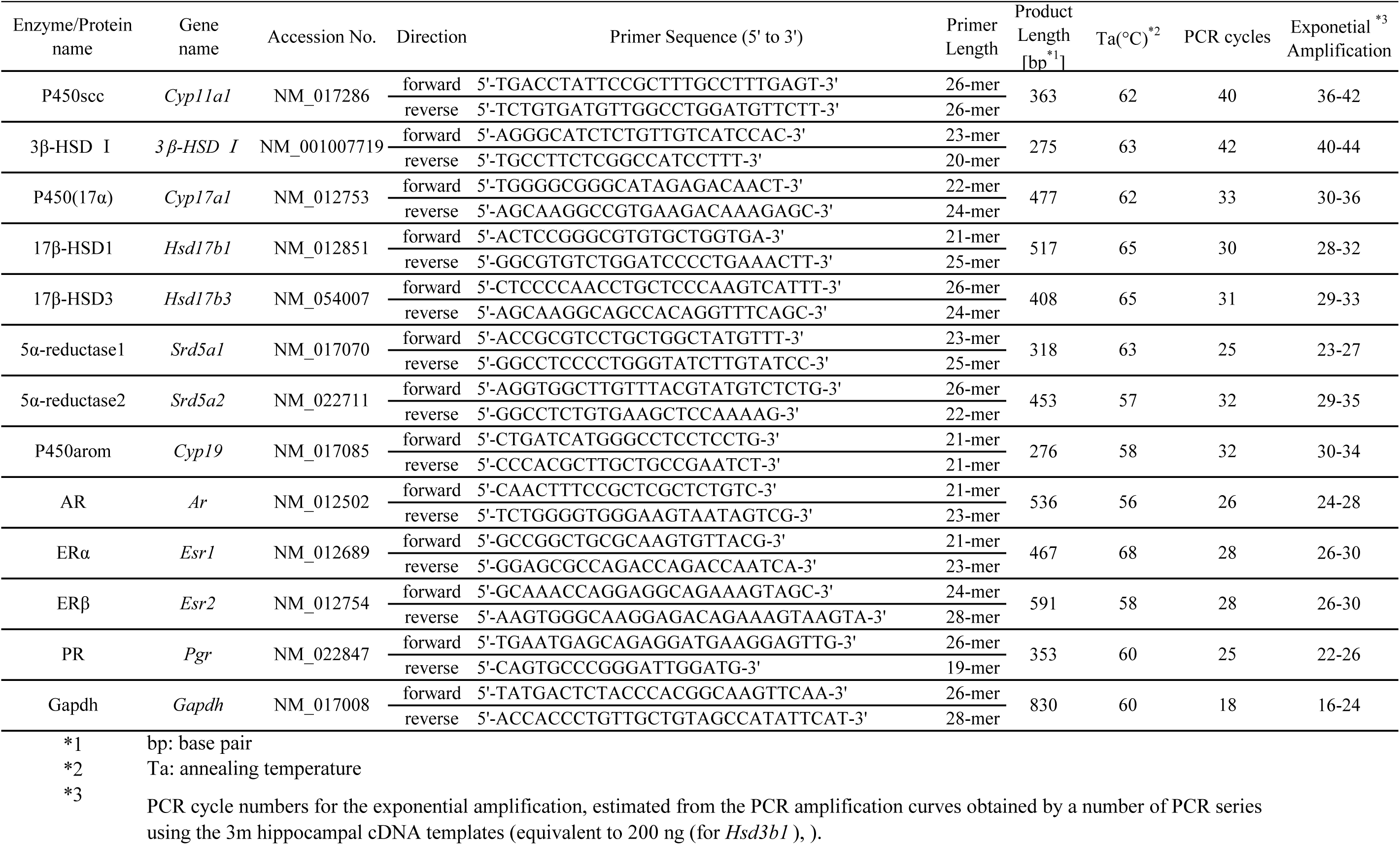
Primers

**Table S2B.**
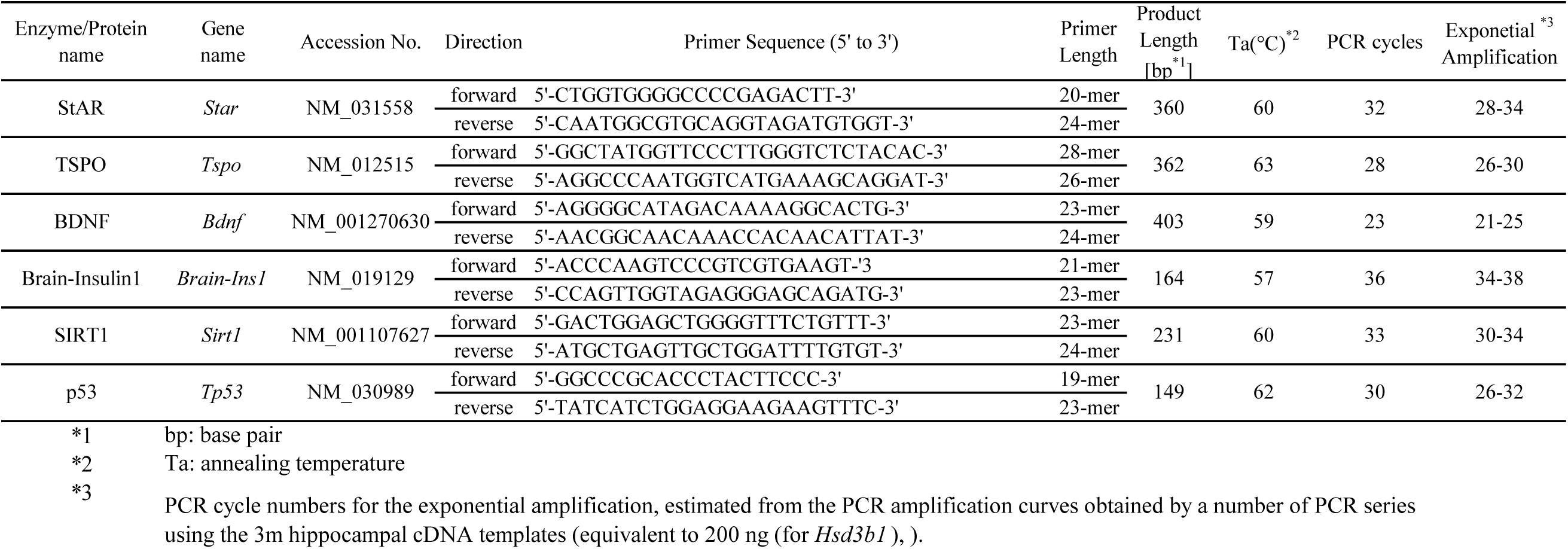
Primers

**Supplementary Table S3.** : Concentrations of T and E2 in the hippocampus at P1, P10, 4 week-old are shown. These concentrations of are significantly lower than those at 3m and 12m. Interestingly, the concentrations of plasma T (0.95 nM) and E2 (0.017 nM) at P10 were significantly lower than hippocampal E2 and T.

|  | P1 | P10 | 4 week-old |
| --- | --- | --- | --- |
| T (nM) | 1.9 | 1.8 | 0.61 |
| E2 (nM) | 2.9 | 0.36 | 0.53 |

